# Retinal neurons establish mosaic patterning by excluding homotypic somata from their dendritic territory

**DOI:** 10.1101/2023.11.17.567616

**Authors:** Christopher Kozlowski, Sarah E. Hadyniak, Jeremy N. Kay

## Abstract

In vertebrate retina, individual neurons of the same type are distributed regularly across the tissue in a pattern known as a mosaic. Establishment of mosaics during development requires cell-cell repulsion among homotypic neurons, but the mechanisms underlying this repulsion remain unknown. Here we show that two mouse retinal cell types, OFF and ON starburst amacrine cells, establish mosaic spacing by using their dendritic arbors to repel neighboring homotypic somata. Using newly-generated transgenic tools and single cell labeling, we identify a transient developmental period when starburst somata receive extensive contacts from neighboring starburst dendrites; these serve to exclude somata from settling within the neighbor’s dendritic territory. Dendrite-soma exclusion is mediated by MEGF10, a cell-surface molecule required for starburst mosaic patterning. Our results implicate dendrite-soma exclusion as a key mechanism underlying starburst mosaic spacing, and suggest that this could be a general mechanism for mosaic patterning across many cell types and species.

## INTRODUCTION

The vertebrate retina contains over 120 cell types, each specialized for particular visual functions (Shekhar and Sanes, 2021). Many of these cell types exhibit mosaic patterning, whereby individual neurons of a given type are distributed into regularly spaced soma arrays (Wässle and Riemann, 1978). This patterning system serves a crucial role in visual processing: It ensures uniform and complete distribution of neural elements across the retina so that the same computations may be performed at all points throughout the visual field. Accordingly, disruption of mosaic patterning impairs retinal circuit function and visually guided behavior (Chen et al., 2013; Ray et al., 2018). Because mosaics are so fundamental to retinal anatomy and functional output, it is critical to understand the precise mechanisms that guide their formation.

The most important developmental phenomenon generating mosaic patterning is local cell-cell repulsion between neurons of the same type (Cook and Chalupa, 2000; Galli-Resta, 2002; Reese and Keeley, 2015). A defining feature of mature mosaics is the “exclusion zone” – i.e., the region surrounding each cell where another homotypic neuron is rarely found (Keeley et al., 2020; Rockhill et al., 2000). Mature mosaics may also exhibit regularity over larger spatial scales; however, modeling studies suggest that local repulsive interactions are sufficient to impose global order upon the entire neuronal array (Eglen et al., 2000; Galli-Resta et al., 1999; Reese and Galli-Resta, 2002). Thus, the key to understanding mosaic patterning is to define the mechanisms by which homotypic neurons repel each other to produce exclusion zones. At present, these mechanisms remain unclear.

Prior work indicates that homotypic repulsion begins shortly after neurogenesis, when newborn neurons have completed their radial migration across the outer neuroblast layer to arrive at their final laminar location (Galli-Resta et al., 1997). At this time, homotypic contacts begin to occur, likely driven by the onset of dendritic outgrowth which coincides with completion of radial migration (Galli-Resta et al., 2002; Reese and Galli-Resta, 2002). These homotypic contacts induce tangential migrations that establish exclusion zones (Amini et al., 2019; Chow et al., 2015; Reese and Galli-Resta, 2002; Reese et al., 1999). However, the anatomical nature of the contacts, and the mechanisms driving cell-cell repulsion once contact has occurred, remain unknown. Cell-surface molecules, such as DSCAM and MEGF10 family members, must be part of the cell-cell repulsion mechanism, as they are needed for homotypic repulsion during mosaic patterning of various cell types (Fuerst et al., 2009; Fuerst et al., 2008; Kay et al., 2012). However, we do not yet know the cellular context in which these molecules act to mediate repulsion, leaving open the question of how repulsion is ultimately accomplished.

Dendritic tiling, a phenomenon whereby neurons establish non-overlapping dendritic territories using homotypic dendritic repulsion (Grueber and Sagasti, 2010), was proposed as a cellular mechanism for carving out exclusion zones (Cook and Chalupa, 2000; Galli-Resta et al., 2002; Huckfeldt et al., 2009). In this model, exclusive dendritic territories could prevent homotypic somata from settling near each other, thereby creating exclusion zones. While dendritic tiling is fairly uncommon in adult retina, a study of mouse horizontal cells showed that dendrites can tile transiently during early development, when regular spacing of cell bodies is being established (Huckfeldt et al., 2009). Based on this finding, transient tiling is now widely considered to be the most likely cellular mechanism underlying exclusion zone formation. Nevertheless, despite its wide acceptance, the transient tiling model has yet to be critically tested. As such, the mechanism by which homotypic neurons avoid each other to generate mosaic patterning remains a long-standing mystery.

Here, to investigate cellular mechanisms underlying homotypic repulsion and exclusion zone formation, we used the cholinergic “starburst” amacrine cells of mouse retina as our model system (Fig. 1A). These cells are advantageous for this purpose not only because they have been a long-standing model for studying mosaics (Galli-Resta et al., 1997; Wässle and Riemann, 1978), but also because they are among the only cell types for which the molecular mechanism driving exclusion zone formation is known (Kay et al., 2012). Starburst neurons comprise two separate interneuron populations that form independent mosaics: the OFF starburst cells of the inner nuclear layer (INL), and the ON starbursts of the ganglion cell layer (GCL). Patterning of both starburst mosaics requires the transmembrane receptor MEGF10, which serves as both a receptor and ligand to mediate homotypic recognition and repulsion (Kay et al., 2012; Ray et al., 2018). In mice lacking *Megf10* gene function, starburst positioning is no longer constrained by the locations of homotypic neighbors; instead, soma positions are random, indicating a complete loss of exclusion zones (Kay et al., 2012). Thus, by understanding the function of MEGF10 at early stages of starburst development, it will be possible to gain insight into the key cellular events that are necessary for neurons to avoid each other during exclusion zone formation.

**Figure 1:**
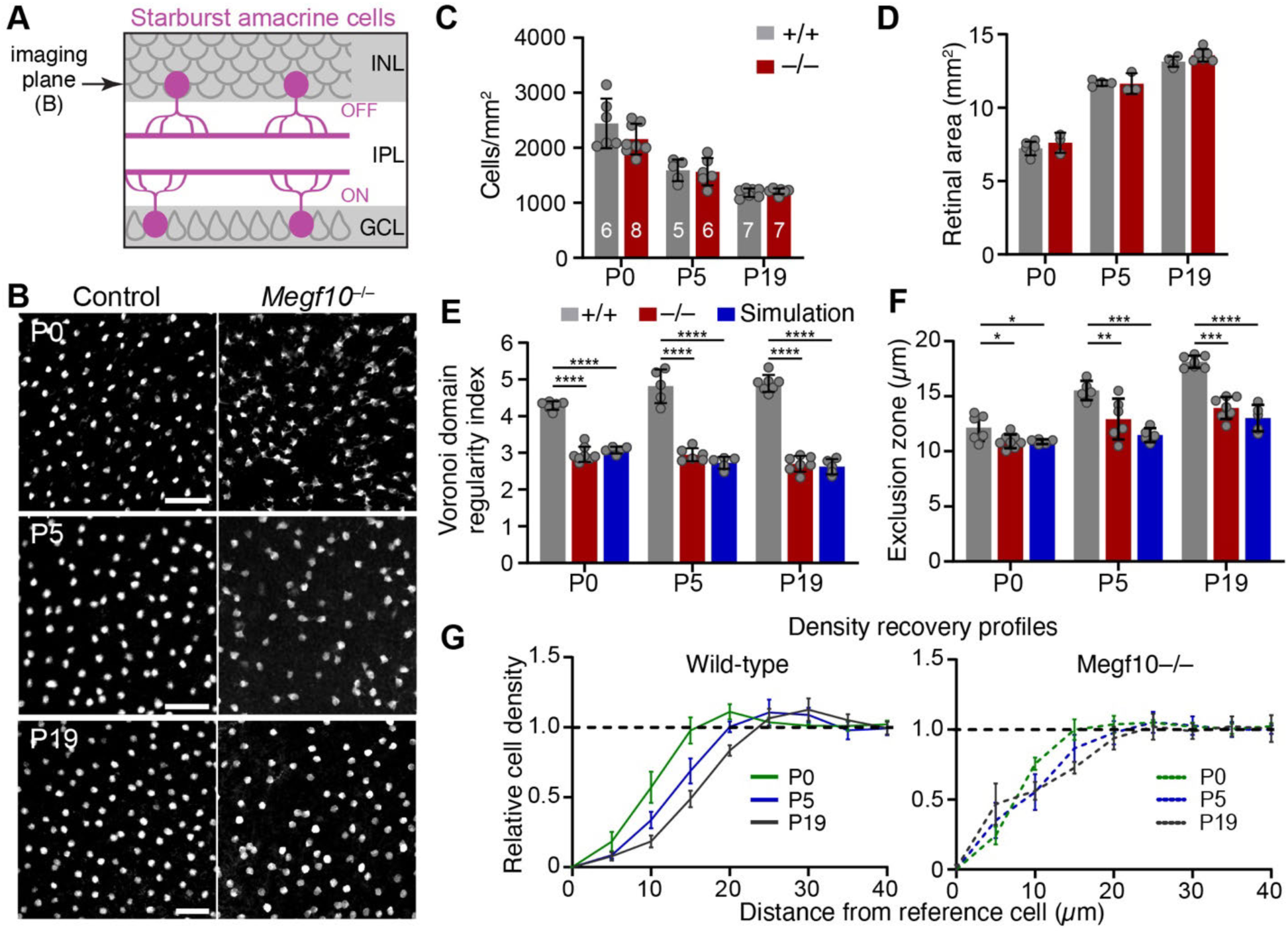
Early establishment of OFF starburst mosaic spacing requires MEGF10. (**A**) Schematic of mature mouse retina, cross-sectional view. Starburst amacrine cell bodies (magenta) are located in inner nuclear layer (INL; OFF starbursts) and ganglion cell layer (GCL; ON starbursts). Starburst dendrites arborize within inner plexiform layer (IPL). Arrow, imaging plane used to acquire en face confocal micrographs (e.g. panel B) for analysis of OFF starburst mosaic patterning. (**B**) Representative images of OFF starburst cell array in *Megf10^−/−^* mutants and wild-type littermate controls (*Megf10^+/+^*). Arrays were labeled at 3 ages using age-appropriate markers: P0, anti-Sox2 (wild-type) or anti-β-galactosidase (mutant), which detects expression of *lacZ* cassette driven by *Megf10* null allele; P5; anti-Sox2; P19, anti-ChAT. (**C,D**) OFF starburst cell density (C) and retinal area (D) were similar in controls and *Megf10* mutants at all ages. Density data (C) is averaged from 3 mid-peripheral images per animal. Sample sizes (numbers of animals) indicated on graph. (**E**) Regularity of OFF starburst array, assessed by VDRI (see Supplemental Fig. S1), in real images and in simulated random arrays matched in cell size and density to real data. Starburst regularity in *Megf10* mutants was significantly lower than in controls, and was indistinguishable from a random distribution. Statistics: 1-way ANOVA (main effects of genotype at each age, p < 1 x 10^−7^) with Tukey’s post-hoc test, ****p < 1 x 10^−6^. P-values for mutant vs. simulated arrays: P0, p = 0.43; P5, p = 0.44; P19, p = 0.82. (**F,G**) Exclusion zone sizes of OFF starbursts (F), measured using density recovery profiles (DRP; see Methods and Supplementary Fig. S1). DRP curves (G,H) show cell density within annuli of increasing size (5 µm increments) around each cell in an imaging dataset. Density in each annulus is normalized to global cell density (dashed line). In wild-type, starburst density at short spatial scales is far lower than global density, indicative of local cell-cell repulsion. Wild-type exclusion zones become larger with time (F; see Results for statistics) as shown by in rightward shift of DRP curve (G). *Megf10* mutants lack bona fide exclusion zones, as their exclusion zone sizes were indistinguishable from random arrays at all ages (F). Statistics (F): 1-way ANOVAs were used to test for genotype effects at each age (main effect of genotype at P0, p = 0.019; at P5, p = 9 x 10^−4^; at P19, p > 1 x 10^−7^) with Tukey’s post-hoc test, *p < 0.05; **p < 0.01; ***p < 0.001; ****p < 1 x 10^−6^. P-values for mutant vs. simulated arrays: P0, p = 0.99; P5, p = 0.19; P19, p = 0.23. Number of mice analyzed for each experimental group (C-G) is indicated in C. Simulations, n = 5. Error bars, SD. Scale bars (B): 50 μm (P0 and P5 are same scale).

To uncover the cellular behaviors controlled by MEGF10 that establish starburst exclusion zones, we built a mouse genetic toolkit for visualizing starburst anatomy at the earliest stages of their differentiation. We discovered that, contrary to the prevailing transient tiling model, starburst dendrites overlap extensively during mosaic formation. Instead, starburst arbors transiently contact the somata of homotypic neighbors, thereby excluding these neighbors from their dendritic territory. In *Megf10* mutants, starburst cells still undertake tangential movements, and their dendrites still contact neighboring starburst somata. However, these contacts are unable to prevent neighboring cells from residing within their arborization territory. Our findings therefore support a new model of mosaic formation whereby dendrite-soma interactions establish exclusion zones. This model applies to both OFF and ON starburst populations; and based on previous anatomical observations, it may generalize as a mosaic patterning mechanism for a wide range of other cell types in both mouse and primate retina (Dacey and Brace, 1992; Wässle et al., 2000).

## RESULTS

### OFF starburst exclusion zones emerge by the day of birth

To investigate the cellular mechanisms by which starburst amacrine cells become patterned into a mosaic, we began by defining the developmental period when exclusion zones first arise. To this end we evaluated development of the OFF starburst mosaic at three developmental timepoints spanning the first three postnatal weeks. We analyzed both wild-type mice and *Megf10* null mutants, which lack OFF starburst exclusion zones at maturity (Kay et al., 2012). Starburst arrays were imaged *en-face* in whole-mount retinas stained for starburst markers Sox2 (Ray et al., 2018; Whitney et al., 2014) or choline acetyltransferase (ChAT; Fig. 1B). Spatial features of the starburst arrays were then analyzed using two standard methods (illustrated in Supplemental Fig. S1). First, regularity of neuronal spacing was measured using the Voronoi domain regularity index (VDRI) (Keeley et al., 2007; Raven et al., 2003); and second, the density recovery profile (DRP) was used to measure exclusion zone size (Rodieck, 1991). The extent of regularity or cell-cell avoidance at each timepoint was determined by comparing real starburst VDRI and DRP data to measurements made on simulated arrays of randomly distributed cells, matched to the cell size and density of the real data (Supplemental Fig. S1). In these random simulations, the only constraint on cell position was that two cells cannot occupy the same physical location. Therefore, for the random arrays, the exclusion zone size is equal to the starburst soma diameter (Kay et al., 2012; Keeley et al., 2020). By contrast, real starbursts – which are not randomly distributed – show larger exclusion zones and more regular cell distributions (i.e., higher VDRI values) than the random simulations (Kay et al., 2012; Whitney et al., 2008) (Fig. 1E,F; Supplemental Fig. S1).

The difference between real and random arrays was used to evaluate developmental changes in OFF starburst mosaic patterning across three timepoints: postnatal day (P)0, P5, and P19. In wild-type mice, exclusion zones and orderly starburst positioning were already present at P0 (Fig. 1E,F). Furthermore, as the retina continued to grow between P0 and P19 (Fig. 1D), exclusion zones became larger (Fig. 1F,G; one-way ANOVA, main effect of age p < 1 x 10–7) and mosaic regularity improved (Fig 1E; one-way ANOVA, main effect of age p = 0.004). Thus, OFF starburst mosaics are refined or sharpened following their initial establishment. By contrast, in *Megf10* mutants, starburst positioning was indistinguishable from random simulations at all ages, and exclusion zones failed to emerge as the retina expanded (Fig. 1E-G). These results were not due to *Megf10* effects on retinal area or cell density, as mutants were comparable to wild-type controls on both measures (Fig. 1C,D). Therefore, MEGF10 is required both for the initial establishment of OFF starburst exclusion zones by P0 as well as the subsequent refinement of starburst mosaic spacing. We conclude that MEGF10-mediated cell-cell interactions driving starburst mosaic development begin prior to P0 and continue into the early postnatal period.

### *Megf10^Cre^* mouse line enables labeling of starbursts during early development

We next sought to determine the anatomical nature of the cell-cell interactions underlying starburst mosaic formation. Because these interactions begin prior to P0 and continue through at least P5 (Fig. 1), this anatomical study requires a tool for labeling nascent starburst dendrites at embryonic and neonatal stages. Previously, we used *Chat^Cre^* mice to drive starburst-specific expression of Cre-dependent membrane-targeted fluorescent proteins (denoted Chat:mGFP; Ray et al., 2018). While this approach is useful for single-cell labeling in neonates, the *Chat^Cre^* driver is not expressed embryonically and yields only sporadic starburst labeling until P5 (Ray et al., 2018; Xu et al., 2016). Therefore, to visualize the complete starburst network at these early ages, we generated a new mouse line expressing Cre in embryonic starburst cells. We targeted the *Megf10* locus for this purpose because, in embryonic retina, newborn starburst cells initiate *Megf10* expression immediately upon completion of radial migration (Kay et al., 2012). This timing coincides with first contact among starburst cells and the earliest emergence of nonrandom spacing (Galli-Resta et al., 1997; Ray et al., 2018). Knock-in mice were generated by inserting a T2A-Cre cassette in place of the endogenous *Megf10* stop codon (Fig. 2A). The targeting construct also contained a FRT-flanked neomycin cassette, used for ES cell selection, within the 3’ untranslated region (UTR) of the *Megf10* transcript. Due to the presence of the neomycin cassette, we refer to the allele derived from this targeting construct as *Megf10^CreNeo^*. Subsequently, we bred *Megf10^CreNeo^*with a germline FLP deleter strain to remove the neomycin cassette, thereby generating a second allele, *Megf10^Cre^* (Fig. 2A).

**Figure 2:**
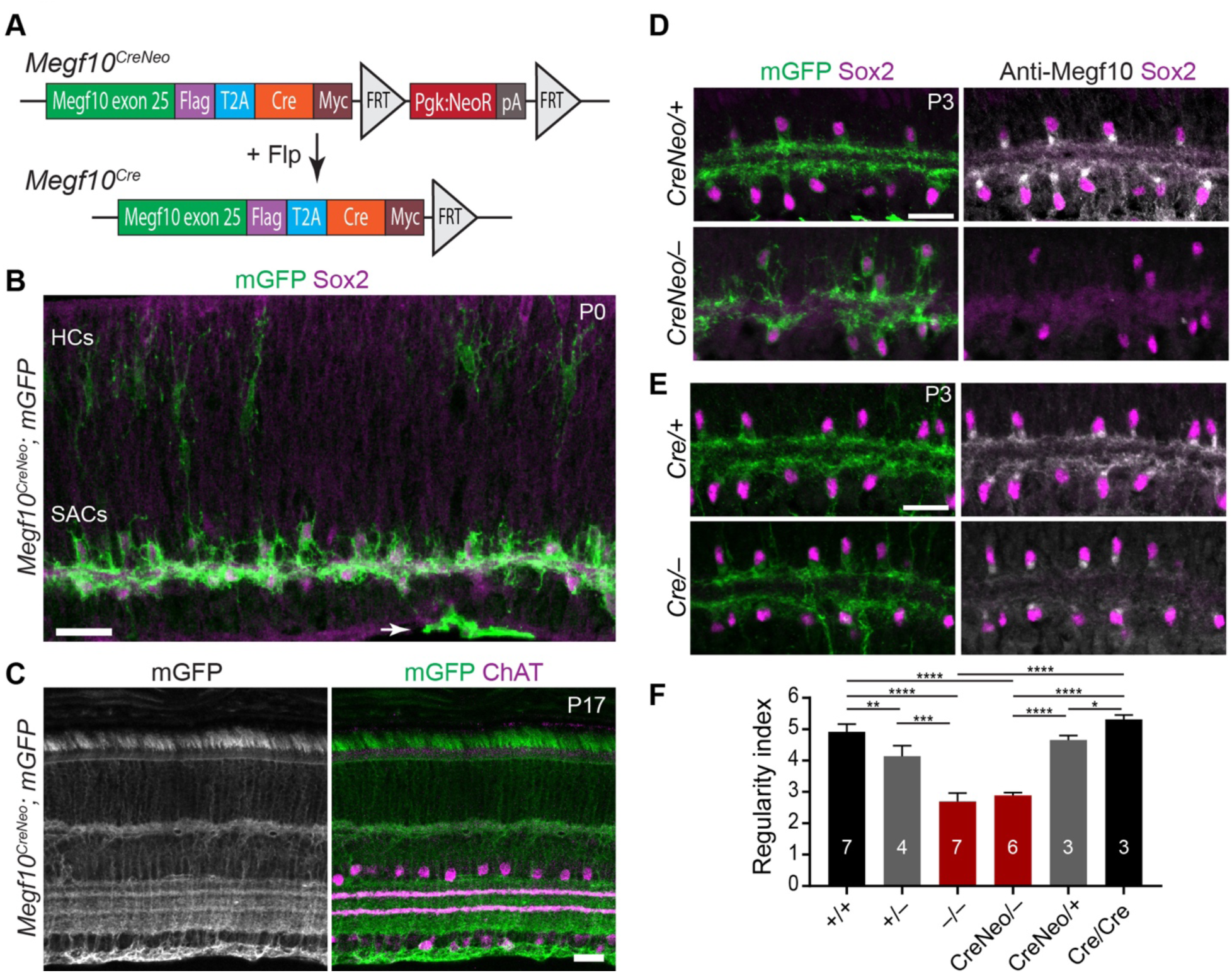
Generation of *Megf10^CreNeo^* and *Megf10^Cre^* mice. (**A**) Schematic of *Megf10* knock-in alleles. Removal of neomycin cassette from *Megf10^CreNeo^*, by breeding with FLP-recombinase germline deleter mouse, generates *Megf10^Cre^* allele. (**B,C**) *Megf10^CreNeo^*expression pattern was assessed by crossing to mGFP Cre reporter line. At P0 (B), most Sox2^+^ starburst amacrine cells (SACs) are GFP^+^, as are horizontal cells (HCs) in outer retina. Occasional GFP-negative Sox2 cells are observed. Sparse labeling is also seen in nerve fiber layer astrocytes (arrow). At P17 (C), Müller glia labeling has become prominent, although ChAT^+^ starburst cells remain GFP^+^. (**D,E**) Comparison of *Megf10^CreNeo^* (D) and *Megf10^Cre^*(E) alleles at P3. Both alleles drive expression of mGFP reporter in most Sox2^+^ starburst cells. The *CreNeo* allele (D) does not support expression of MEGF10 protein, as *CreNeo/–* mice lack MEGF10 immunoreactivity and show IPL innervation defects typical of *Megf10^−/−^* animals (Ray et al., 2018). *Cre/–* animals have normal starburst anatomy and still express MEGF10 protein (E). (**F**) Mosaic regularity of OFF starburst cell arrays, measured by Voronoi domain regularity index, in mice carrying various *Megf10* alleles. The *CreNeo* allele behaves similarly to the null (–) allele, causing mosaic defects, whereas the *Cre* allele behaves similarly to the wild-type (+) allele. Statistics: One-way ANOVA with Tukey’s post-hoc test. Colors indicate pairs of genotypes that are not significantly different from each other (*+/+* vs. *Cre/Cre*, p = 0.17; *+/–* vs. *CreNeo/+*, p = 0.58; *–/–* vs. *CreNeo/CreNeo*, p = 0.74), but which are significantly different from the other color groups. *p = 0.018; **p = 0.0003; ***p < 1 x 10^−6^; ****p < 1 x 10^−7^. Sample sizes: n = 6 animals (+/+; CreNeo/–) or n = 4 (others). Error bars, SD. Scale bars = 25 μm.

To validate that *Megf10^CreNeo^* and *Megf10^Cre^*lines faithfully recapitulate endogenous *Megf10* expression patterns, we characterized Cre activity at three timepoints – P0, P3, and P17 – for which the *Megf10* expression pattern has been thoroughly documented (Kay et al., 2012; Ray et al., 2018). To this end, each line was crossed to a Cre-dependent membrane-targeted GFP (mGFP) reporter (Muzumdar et al., 2007). At P0 and P3, *Megf10*-driven mGFP labeling (denoted Megf10:mGFP) was observed in the vast majority of ON and OFF starburst cells as well as horizontal cells (Fig. 2B,D). Astrocytes in the nerve fiber layer were also GFP^+^ (Fig. 2B). Later, by P17, Müller glia also became labeled (Fig. 2C). Each of these findings is consistent with our previous analysis of *Megf10* mRNA and protein expression (Kay et al., 2012; Ray et al., 2018). To determine how the knock-in strategy affects *Megf10* gene function, *CreNeo* and *Cre* mutant mice were stained with antibodies to MEGF10. We only detected MEGF10 protein when the neomycin cassette was removed from the 3’ UTR (Fig. 2D,E), suggesting that *Megf10^CreNeo^*is likely to be a strong loss-of-function or null allele. Consistent with this interpretation, *Megf10^CreNeo/–^* mice phenocopied starburst dendritic and soma positioning errors seen in *Megf10^−/−^*null mutants (Fig. 2D,F; Ray et al., 2018). By contrast, *Megf10^Cre/–^* mice retained expression of MEGF10 protein and did not exhibit starburst phenotypes (Fig. 2E,F). Altogether, these results show that the two *Megf10*-driven Cre lines serve as reliable early markers of starburst cells and can be deployed to preserve (*Megf10^Cre^*) or abrogate (*Megf10^CreNeo^*) MEGF10 function depending on experimental needs.

### Starburst dendrites contact neighboring starburst somata during mosaic formation

Using these new mouse lines, we investigated the anatomy of starburst cell-cell contacts during the late prenatal and early postnatal period when mosaic spacing and exclusion zones are established. As a prenatal timepoint we chose embryonic day (E)16, when nonrandom cellular distributions are first arising and when newly arrived post-migratory starbursts are still being incorporated into the nascent mosaic (Galli-Resta et al., 1997; Ray et al., 2018). The prevailing “transient tiling” model of mosaic formation predicts that starbursts should tile their dendrites during this period, even though they do not tile in adulthood (Fig. 3A,B) (Cook and Chalupa, 2000; Huckfeldt et al., 2009; Keeley et al., 2007; Reese and Galli-Resta, 2002). To investigate this possibility we labeled the full starburst population with Megf10:mGFP (Fig. 3D-G), as well as labeling single starbursts with Chat:mGFP (Fig. 3H-K). This analysis revealed that, starting by E16 (Fig. 3D-F) and continuing through P1 (Fig. 3D,G-K), starburst anatomy was inconsistent with tiling: Rather than stopping at their neighbor’s dendritic tips, starburst arbors frequently extended all the way to neighboring starburst cell bodies (see schematic, Fig. 3C). This behavior was unchanged in *Megf10* mutants (Fig. 3E). Importantly, the propensity to contact neighboring somata was observed even for the most immature E16 starbursts – those that still had the radial morphology of a migrating cell and had not yet begun to innervate the IPL (Fig. 3F). This observation suggests that starburst dendrites can directly target somata without first moving through a phase of transient tiling.

**Figure 3:**
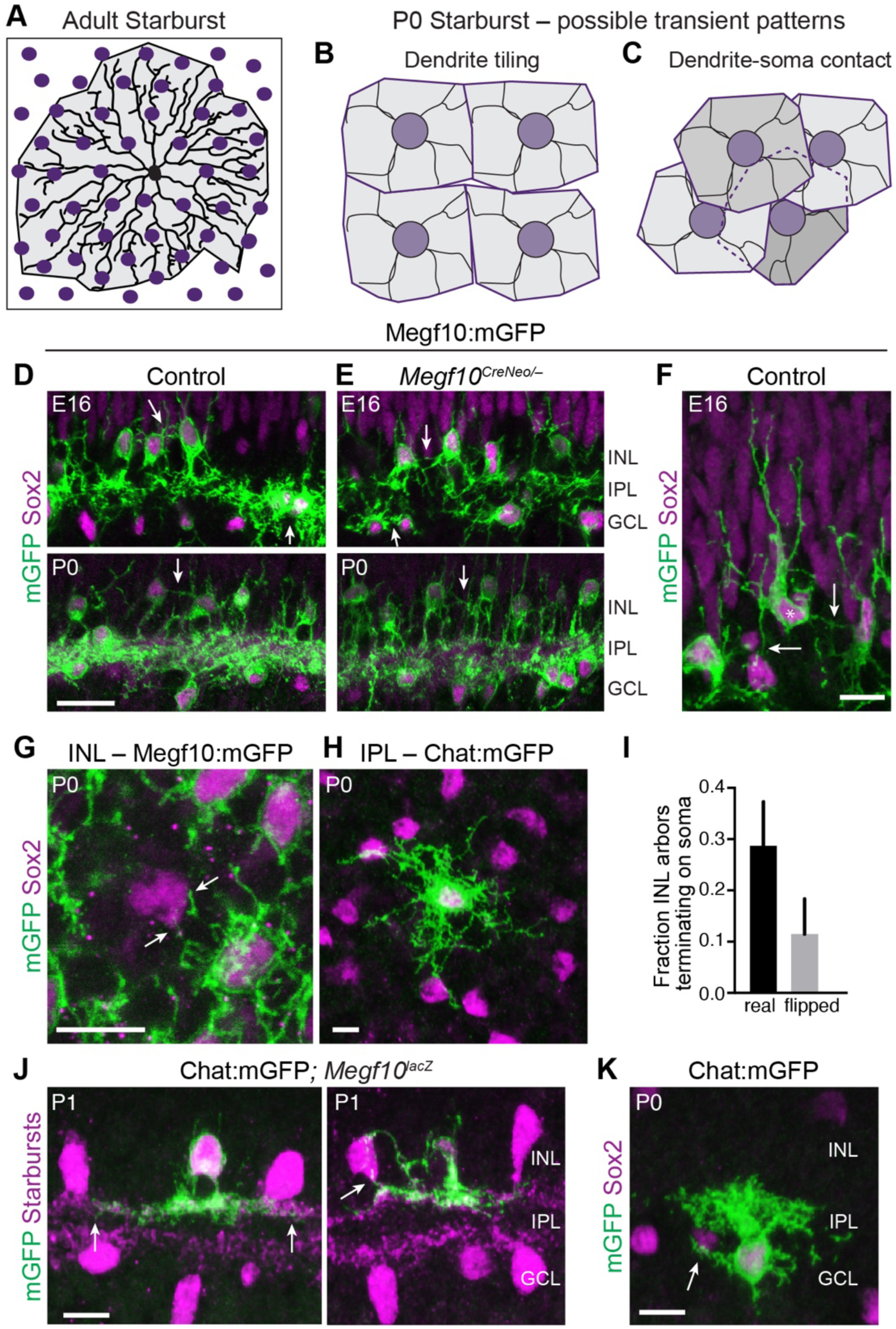
Nascent starburst dendrites contact neighboring starburst somata. (**A**) Schematic showing morphology of a single starburst neuron (black) in adult retina, and relationship of its dendrite territory (gray shading) to neighboring starburst cell bodies (purple). In adults, starburst dendrites extend well beyond the first ring of adjacent homotypic neighbors. (**B,C**) Schematic illustrating two possible models for establishment of exclusion zones by neonatal starburst cells. The predominant tiling model (B) holds that starburst cells (purple) transiently establish nonoverlapping dendritic territories (gray shading) to generate local cell-cell avoidance. (C) Avoidance could also emerge from transient repulsive interactions between dendrites and neighboring starburst somata (**D,E**) Megf10:mGFP labeling at E16 (top) and P0 (bottom). Sox2^+^mGFP^+^ starburst cells send dendrites into IPL, but also project arbors into soma layers (INL, GCL). Soma layer arbors contact neighboring starburst cell bodies (arrows), both in controls (D) and *Megf10* mutants (E). Note that, at E16, Sox2 also marks both progenitor cells in the outer neuroblast layer. However, within INL and GCL, starbursts are the only postmitotic neurons that express Sox2. (**F**) Example of radially migrating starburst cell at E16. While still migrating, this cell projects arbors that contact adjacent Megf10^+^Sox2^+^ starburst somata that have already completed their migration to INL (arrows). (**G,H**) Dendro-somatic contact between OFF starbursts at P0, imaged en face. G, INL arbors labeled by Megf10:mGFP. A rare GFP-negative Sox2^+^ starburst cell (center) is contacted by GFP^+^ arbors (arrows). H, Chat:mGFP-labeled OFF starburst cell, imaged at IPL level and overlaid onto INL Sox2 array. IPL dendrites extend as far as the next ring of Sox2^+^ cell bodies. (**I**) Quantification of INL soma contacts, using Chat:mGFP images similar to H. Arbor tips were more likely to terminate on a starburst somata in real images as compared with control images in which one channel was flipped about the X and Y axes. Sample size, 122 arbors from 22 cells. Error bars, 95% confidence intervals. (**J,K**) Cross-sectional view of dendro-somatic contacts between OFF (J) and ON (K) starburst cells. Chat:mGFP was used for single-cell labeling; full starburst population labeled by *lacZ* transgene driven from *Megf10* locus (J) or anti-Sox2 (K). GFP^+^ arbors within IPL and within soma layers typically extend only as far as neighboring starburst somata. In some cases dendrites appear to grab neighboring cells (J, right; K). Images in J (right-hand panel only) and K are reproduced from Ray et al., 2018. Scale bars: 25 µm (A) or 10 µm (all other panels).

Several aspects of starburst anatomy during the E16-P1 period suggested that starburst dendrites selectively target neighboring somata rather than touching them by chance. First, consistent with our prior studies (Ray et al., 2018), we found that starbursts possess transient dendritic arbors that are specialized for soma contact. In addition to their main dendritic arbor, which ramifies within the inner plexiform layer (IPL; Fig. 1A), starbursts also produce a transient arborization within their cell body layer – i.e. the INL or GCL – that is eliminated by P3 (Fig. 3DE,G; compare to Fig. 2D,E). These transient soma layer projections are a prominent source of dendrite-soma contacts (Fig. 3D-G). Using Chat:mGFP to label single neurons, we found that 73% of P0-1 OFF cells contacted a neighboring starburst soma using INL-directed arbors (n = 16/22). Furthermore, the frequency of starburst dendrite-soma contact within the INL was greater than the rate expected by chance (Fig. 3I), supporting the idea that such contacts are selective. Starburst IPL arborizations also showed evidence of selective soma contact at P0-1. Alignment between dendrite tips in the IPL and neighboring somata was remarkably precise (Fig. 3H,J,K); contacts were typically localized to the base of the starburst cell where it touched the IPL, but in some cases dendrites even branched out of the IPL to contact INL cell bodies (Fig. 3J). Together these findings indicate that, instead of tiling (Fig. 3B), starburst dendrites are transiently recruited to target neighboring homotypic somata during the perinatal period (Fig. 3C). This observation raises the possibility that dendro-somatic contacts could play a role in establishing starburst mosaic spacing.

### OFF starburst dendrites exclude neighboring somata from their territories

We next sought to determine whether starburst dendro-somatic contacts contribute to establishment of mosaic spacing. Based on the observed anatomy during the perinatal period (Fig. 3C), a plausible hypothesis is that dendritic contacts establish exclusion zones by transiently restricting starburst somata from entering another cell’s dendritic territory. If this model is correct, we can make the following two predictions: 1) During mosaic formation, starburst arbors should demarcate a zone within which neighboring cells are rarely found; and 2) this dendro-somatic avoidance behavior should be impaired in *Megf10* mutants, which lack starburst exclusion zones (Fig. 1F).

To test the first prediction, we marked individual OFF starburst cells using Chat:mGFP and measured how many starburst somata were enclosed within the reference cell’s dendritic territory. For each cell, measurements were made on the real image (n = 36) as well as a family of “unmatched” images (n = 32 per cell) in which the reference cell dendritic outline was placed arbitrarily onto OFF starburst arrays from different retinal locations (Fig. 4A). This unmatched control condition was used to quantify the number of enclosed cells expected by chance in the absence of spatial coordination between dendrites and neighboring somata. At P0, the majority of real starburst arbors contained only a single cell body – the soma belonging to the reference Chat:mGFP^+^ cell (Fig. 4D). By contrast, unmatched simulations showed significantly higher frequencies of multiple cell enclosure (Fig. 4D) and enclosed more neighboring cells, on average, than real arbors (Fig. 4C). This tendency for real arbors to enclose fewer neighbors was observed not only when we compared the entire real and simulated datasets (Fig. 4C), but also when we performed a more stringent test in which each real arbor was compared only to simulations generated from that same arbor (Fig. 4F). Altogether, this analysis shows that neighboring starburst somata are found within the reference cell arbor less often than expected by chance. This finding supports the conclusion that P0 starbursts avoid residing within neighboring cells’ dendritic territories (Fig. 4G).

**Figure 4:**
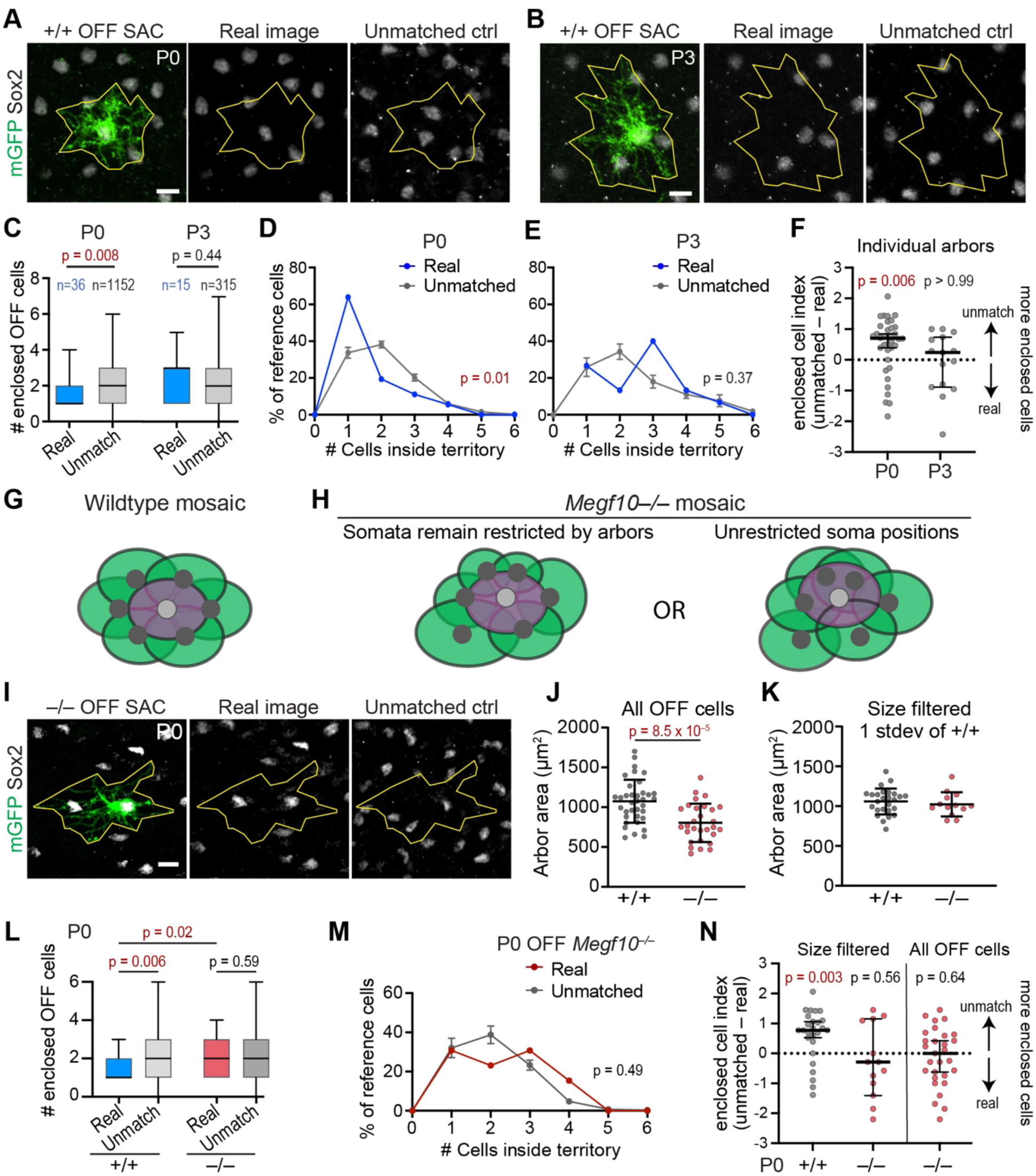
OFF starburst dendritic arbors exclude homotypic somata in a MEGF10-dependent manner. (**A,B**) Representative Chat:mGFP^+^ OFF starburst reference cell from P0 (A) or P3 (B) wild-type (+/+) retina, imaged en face. Anti-Sox2 marks neighboring starburst amacrine cells (SACs). Real images (left, center) show relationship of Sox2 array to reference cell dendritic territory (yellow polygon). In unmatched control (“ctrl”) image, the polygon (yellow) was placed onto the OFF starburst array from a different image. (**C**) Box and whisker plot summarizing number of OFF Sox2^+^ cells fully enclosed by real dendritic territory polygons (blue) or unmatched controls (gray). Box, interquartile range; black line, median. Sample sizes: P0, n = 36 real arbors from 7 animals; n = 32 unmatched images per real arbor (1,152 total control images). P3, n = 15 real arbors from 3 mice; n = 21 unmatched images per arbor (315 total controls). Statistics, two-tailed Mann-Whitney test. (**D,E**) Histograms showing fraction of reference cells enclosing *n* somata at P0 (D) or P3 (E). Unmatched control images enclosing zero cells were excluded, so as to match real data where at least one cell (the reference cell) was inside each arbor. Sample sizes: real arbors as in C; unmatched P0, n = 1,117; unmatched P3, n = 297. Statistics: chi-squared test. (**F**) For each reference cell, an “enclosed cell index” was calculated by subtracting the real Sox2 count from the mean number counted in unmatched controls generated using that same arbor (number of controls per arbor as in C). If real and unmatched counts are equal (i.e. no difference between groups), this index will be equal to 0 (dashed line). Deviations above or below dashed line indicate more cells in unmatched or real conditions respectively. At P0, there were significantly more cells within unmatched arbors, showing that real arbors contain fewer neighboring cells than expected by chance. No such difference was detectable at P3. Statistics: Wilcoxon one-sample test with theoretical median of 0. (**G,H**) Schematics illustrating how loss of *Megf10* might alter soma-dendrite arrangements. In wild-type (G), somata avoid neighboring dendritic territories, establishing exclusion zones and regular spacing. In *Megf10* mutants (H), cells are randomly positioned. Two possible explanations for loss of exclusion zones are illustrated. H, left: Cell position remains restricted by arbor exclusion, in which case random positioning must result from variability in arbor sizes (green territories). H, right: Mutant somata are no longer excluded from neighboring arbor territories. (**I**) Representative Chat:mGFP reference cell from P0 *Megf10^−/−^* retina, shown with its real starburst array (center) and placed onto an unmatched OFF starburst array (right). (**J**) Dendritic arbor areas in mutant P0 OFF starbursts (n = 31 arbors from 7 *Megf10^−/−^* animals) are smaller than littermate controls (wild-type sample size: n = 36 as in B). Statistics, two tailed t-test. (**K**) Filtering of arbors >1 standard deviation (stdev) from wild-type mean generates a reference cell dataset in which arbor sizes are comparable between genotypes (wild-type arbors n = 26; *Megf10* mutant arbors n = 13). (**L**) : Box and whisker plot (as in C) quantifying Sox2^+^ cell enclosure in P0 *Megf10* mutants and wild-type controls using the size-matched dataset (K). Mutant arbors enclosed more cells than wild-type arbors but a similar number as unmatched controls, consistent with a loss of dendrite-soma exclusion in mutants. Sample sizes: real arbors as in K; unmatched images: wild-type, n = 32 per arbor, n = 832 total; mutant, n = 20 per arbor, n = 260 total. Statistics as in C. (**M**) Histogram of enclosure frequencies (as in D,E) using *Megf10* mutant arbors. Histogram for real mutant arbors (sample size as in K) was indistinguishable from chance rate measured using unmatched images (n = 245; unmatched images enclosing zero cells were excluded). Statistics as in D,E. (**N**) Enclosed cell indices for individual P0 arbors were plotted (as in F) for the size-matched dataset (left) and the full dataset of mutant OFF reference cells (right; see F for full wild-type dataset). Wild-type arbors contained fewer Sox2 somata than expected by chance (i.e., unmatched condition), but mutant arbors enclosed a similar number of neighboring cells in both real and unmatched conditions. Sample sizes: Left, as in K,M; right, n = 29 real arbors, 20 unmatched images per arbor (zero values excluded, n = 479 total simulations). Error bars, mean ± SD (J,K); mean ± SEM (D,E,M) median ± 95% CI (F,N); min-max values (C,L). P-values shown on graphs; red text denotes value below alpha threshold. Scale bars, 10 µm. Also see Supplemental Figure S2.

We next asked how long this exclusionary behavior persists. By P3, dendrites have grown larger (Supplemental Fig. S2A) and extend past the first ring of adjacent homotypic cells. We therefore surmised that the relationship between arbor territories and soma positioning would likely be different than at P0. Indeed, at P3 there was no difference in soma enclosure frequency between real and unmatched arbors, demonstrating that dendrites no longer exclude neighboring somata (Fig. 4B-F). Together these findings indicate that starbursts can exclude homotypic somata from their dendritic territories, and that this behavior is limited to the developmental time when exclusion zones first arise.

### MEGF10 enforces exclusion between OFF starburst dendrites and neighboring somata

To learn whether the exclusionary relationship between dendrites and somata is relevant for starburst mosaic formation, we tested the extent to which exclusion persists in *Megf10* mutants. If the random positioning of *Megf10^−/−^* OFF starbursts at P0 (Fig. 1) is caused by failure of soma-dendrite exclusion, we would expect more mutant starbursts to be located within the reference cell dendritic territory than in wild-type animals (Fig. 4H, right panel). However, it is also possible that random positioning is caused by another mechanism, in which case we expect dendrite-soma exclusion to be unaffected in mutants (Fig. 4H, left panel). To distinguish these possibilities, we began by assessing the frequency of neighboring soma enclosure by *Megf10^−/−^* Chat:mGFP^+^ dendrites. We found that real *Megf10^−/−^* GFP^+^ dendrites enclosed starburst cell bodies at a similar frequency to randomized unmatched controls (Fig. 4N; Supplemental Fig. S2C), suggesting that, in contrast to wild-type, mutant somata are not excluded from dendrite territories. However, we also noted that dendritic arbors of *Megf10^−/−^* starbursts were smaller than wild-type arbors (Fig. 4J), complicating efforts to compare enclosure rates between genotypes. To control for arbor size effects, we filtered both wild-type and mutant reference cell datasets to include only those arbors that were within 1 standard deviation of the wild-type mean size. In this way, we obtained a set of wild-type and mutant reference cells with comparable arbor sizes (Fig. 4K). Using these size-matched datasets we found that dendrite-soma exclusion was impaired in *Megf10* mutants, as mutant arbors enclosed significantly more somata than wild-type arbors (Fig. 4L). Moreover, the frequency of cell enclosure by real mutant dendrites was indistinguishable from unmatched controls, suggesting that mutant arbors cannot influence neighboring cell locations (Fig. 4L-N).

To bolster evidence for this soma-exclusion model, we also examined an alternative model of the *Megf10* mutant phenotype in which soma exclusion is unaffected. In order for soma positions to become random while still preserving dendrite-soma exclusion, the sizes of mutant arbors must necessarily become more variable than wild-type arbors (see illustration, Fig. 4H). However, contrary to this prediction, we did not detect differences in arbor size variability between wild-type and mutant starbursts (Levene’s variance test, f-ratio = 0.45, p = 0.51; note similarity of standard deviations in Fig. 4J). This finding strongly suggests that the alternative model is not plausible, thereby providing further support for our conclusion that failure of dendro-somatic exclusion underlies the *Megf10* phenotype. Altogether, this series of experiments indicates that *Megf10* is required for OFF starburst dendrites to exclude neighboring somata, implicating this mechanism in the establishment of exclusion zones required for mosaic spacing.

### Starburst tangential movements occur normally in *Megf10^−/−^* retina

We next investigated why *Megf10* mutant arbors are unable to exclude neighboring somata. In wild-type retina, starburst cells undertake lateral movements within the tangential plane of the retina; these serve to reposition neurons from the site of their birth, which is random, into an orderly mosaic array (Amini et al., 2019; Chow et al., 2015; Galli-Resta et al., 1997; Reese and Galli-Resta, 2002; Reese et al., 1999). Our results so far (Fig. 4) suggest that these tangential movements are normally constrained by contact with neighboring dendritic arbors, whereas in *Megf10* mutants, dendritic contacts do not lead to uniform spacing of starburst cell bodies. One possible explanation for this phenotype is that mutant somata are unable to move from the site of their birth; in this case, repulsive contacts with neighboring dendrites would not have an opportunity to influence soma position. Alternatively, mutant starbursts may still move but their movements are not properly constrained by dendrite contact, such that they are able to enter neighboring arbor territories.

To distinguish between these two models, we tested whether *Megf10^−/−^*starbursts are capable of tangential movements. Using a well-established approach used in previous studies (Reese et al., 1995; Reese et al., 1999), the extent of tangential dispersion was evaluated by marking clones of cells derived from a small subset of retinal progenitors, and then measuring the lateral displacement of starburst progeny from their clone of origin (Fig. 5A). To mark progenitors we used a Pax2-Cre BAC transgenic line (Ohyama and Groves, 2004), which we found to be expressed by a small population of progenitors distributed across the retina (Fig. 5A). When crossed to a tdTomato Cre-reporter mouse, Pax2-Cre drives reporter expression in these progenitors and their clonally labeled progeny (Fig. 5A). In control P2 retinas, this labeling revealed densely packed radial columns as well as laterally displaced Tomato^+^ cells, which have moved away from their clone of origin via tangential migration. Some Sox2^+^ starburst neurons were found among the laterally displaced Tomato^+^ group, while other Tomato^+^ starbursts remained aligned with the labeled clones, indicating that they had not migrated tangentially (Fig. 5A). Quantifying the fraction of starbursts in each group, we found that about half of all control starbursts had migrated tangentially to settle outside of clones (Fig. 5B). This migration frequency is consistent with previous studies examining starburst tangential dispersion using different transgenic mice (Reese and Galli-Resta, 2002; Reese et al., 1995). In *Megf10* mutants, starburst cells still dispersed tangentially away from their clone of origin. Indeed, the frequency of tangential dispersion was unchanged between mutant and control retinas (Fig. 5A,B). Moreover, there was no difference in the average displacement distance for starbursts located outside of clones (Fig. 5C,D). These findings indicate that starburst tangential motility is unaffected by loss of *Megf10*, implying that the mutant phenotype arises because neighboring dendrites are unable to prevent motile starbursts from moving into their arbor territories.

**Figure 5:**
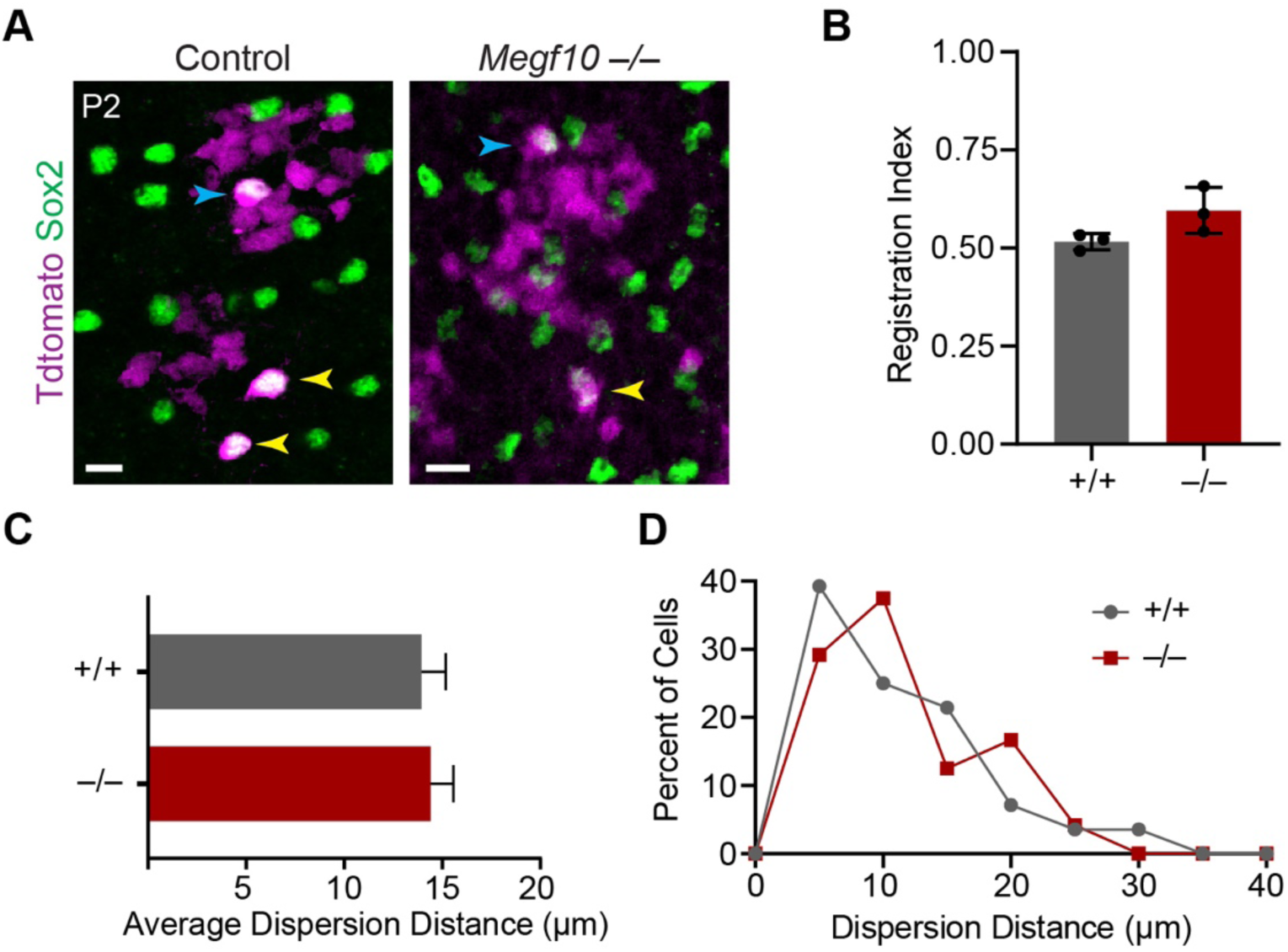
Starburst lateral movements are normal in *Megf10* mutants. (**A**) En face views of INL in P2 retinal wholemounts from Pax2-cre mice carrying tdTomato Cre reporter. Cre drives expression of tdTomato in columns of neurons derived from the same progenitor cell. Sox2^+^ starburst cells are detected both within clonal columns (blue arrows), and also outside of the columns in cases where starbursts have undergone tangential movements away from their clone of origin (yellow arrows). OFF starbursts move laterally both in *Megf10^+/+^* wild-type controls (left) and *Megf10^−/−^* mutants (right). Scale bar, 10µm. (**B**) Migration frequency measured using registration index, quantified as fraction of tdTomato^+^ starbursts remaining aligned within clonal columns (i.e. cells that failed to undergo tangential migration). In both wild-type controls and mutants, about half of starbursts move tangentially away from their clone of origin. Sample sizes: n = 3 animals per genotype, 50-60 cells per animal (wild-type, 196 cells total; mutant, 163 cells total). Statistics, two-tailed t-test, p = 0.09. (**C,D**) Dispersion distance was quantified for starburst cells that were outside their original clonal column (n = 24 wild-type, n = 28 mutant). Average distance from starburst cells to edge of nearest column was unchanged in *Megf10* mutants (C, two-tailed t-test, p = 0.78). The distribution of dispersion distances across the OFF starburst population was also unchanged in mutants (D). Error bars, SD.

### Dendrite-soma exclusion also patterns the ON starburst mosaic

To learn whether the mechanisms that generate the OFF starburst mosaic are applicable to other cell types, we next investigated development of the ON starburst mosaic. We began by examining developmental timing of ON starburst mosaic patterning (Fig. 6). Similar to OFF cells (Fig. 1), the ON starburst mosaic was already present at P0 and continued to refine over the postnatal period (Fig. 6C-E). By contrast, in *Megf10* mutants, the ON array lacked exclusion zones at P0 and regularity failed to improve with age (Fig. 6C). To test whether dendrite-soma exclusion contributes to ON starburst patterning, we analyzed the spatial relationship between single Chat:mGFP^+^ arbors and neighboring ON somata, using the same unmatched randomized control methodology to measure the chance rate of enclosure (Fig. 7A). At P0, real ON starburst arbors contained fewer neighboring somata than expected by chance, with the majority (>60%) of arbors enclosing only the reference Chat:mGFP^+^ soma (Fig. 7A-C,J). This exclusionary relationship between dendritic arbors and neighboring somata was lost in *Megf10* mutants (Fig. 7G-L). Together, these results indicate that both ON and OFF starburst mosaics develop similarly: In both cases, dendrites repel neighboring somata in a *Megf10*-dependent manner to establish exclusion zones.

**Figure 6:**
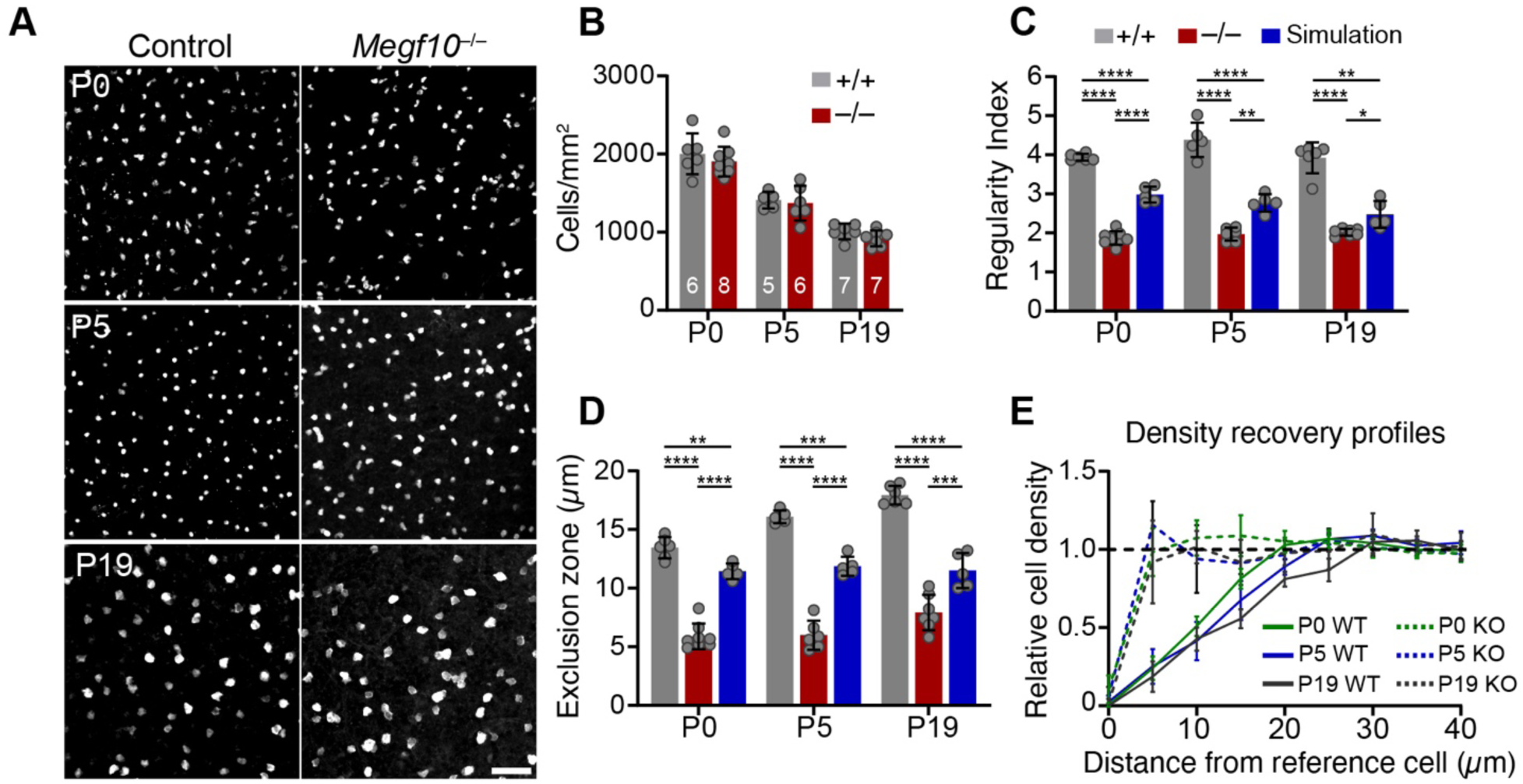
Early establishment of ON starburst mosaic spacing also requires MEGF10. (**A**) Representative images of ON starburst array across development in wild-type controls and *Megf10* mutants. Cells were labeled with anti-Sox2 (P0, P5) or anti-ChAT (P17). (**B**) ON starburst cell density did not differ between *Megf10* mutants and littermate controls (two-way ANOVA, no main effect of genotype, p = 0.18). (**C**) ON starburst regularity, assessed by VDRI, was lower in mutants than in controls. Mutant array regularity was also lower than for simulated random arrays. Statistics: 1-way ANOVA (main effect at all ages, p > 1 x 10^−7^) followed by Tukey’s post-hoc test, *p < 0.05, **p < 0.01, ****p < 1 x 10^−5^. (**D,E**) Exclusion zone size (D), measured using the DRP (E), increased over time in wild-type control mice (one-way ANOVA to test for age effects in wild-type group only, main effect of age p = 4 x 10^−7^). In mutants, exclusion zones were smaller than in controls or in simulated data. Note leftward shift of DRP curves in mutants indicating mild cell aggregation (E). Statistics (D): 1-way ANOVAs were used to test for genotype effects at each age (main effect of genotype at all ages, p > 1 x 10^−7^) with Tukey’s post-hoc test, **p < 0.01; ***p < 0.001; ****p < 1 x 10^−5^. Error bars, SD. Scale bars, 50 µm. Also see Supplemental Fig. S2.

**Figure 7:**
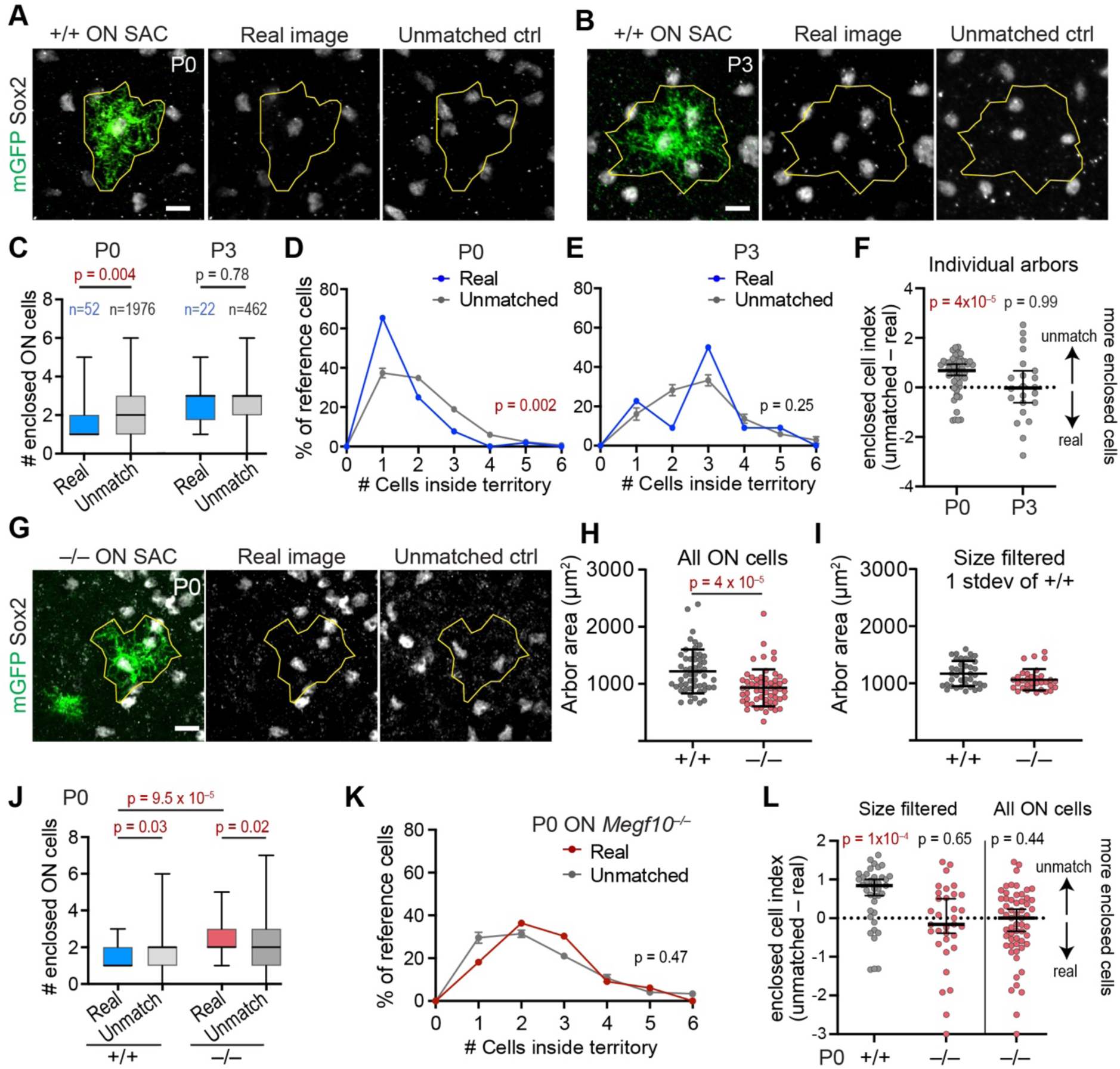
ON starburst dendritic arbors exclude homotypic somata. (**A,B**) Representative Chat:mGFP^+^ ON starburst reference cell from P0 (A) or P3 (B) wild-type (+/+) retina. Sox2 marks neighboring starburst amacrine cell (SAC) somata. Left, center: real images; right, unmatched control condition with dendritic polygon (yellow) moved arbitrarily to a new ON starburst image. (C) Box and whisker plot summarizing number of ON Sox2^+^ cells fully enclosed by real dendritic territory polygons (blue) or unmatched controls (gray). Box, interquartile range; black line, median. Sample sizes, P0, n = 52 real GFP^+^ cells from 7 animals; n = 38 unmatched images per arbor, n = 1,976 total controls; P3, n = 22 real arbors; n = 21 unmatched images per arbor, n = 462 total controls. Statistics, two-tailed Mann-Whitney test. (**D,E**) Histograms showing fraction of ON reference cells enclosing *n* somata at P0 (D) or P3 (**E**). Sample sizes: real arbors as in C; unmatched controls (zero values excluded), P0, n = 1,827; P3, n = 457. Statistics, chi-squared test. (**F**) The enclosed cell index (unmatched arbor mean Sox2 counts – real arbor Sox2 counts) plotted for each individual arbor. Dashed line at zero represents expected value if there is no difference between real and unmatched groups. At P0 but not P3, real arbors systematically enclosed fewer neighboring Sox2 cells than unmatched arbors. Statistics, Wilcoxon one-sample test with theoretical median of 0. (**G**) Representative images of ON starburst dendritic arbors in *Megf10* mutants, together with the ON starburst soma array (Sox2). Real images, (left, center); control unmatched images (right). (**H**) Average ON starburst arbor size is smaller in *Megf10* mutants than in controls (wild-type, n = 52 as in B; mutant, n = 58 cells from 7 animals; two tailed t-test. (**I**) Filtering of arbors >1 standard deviation (stdev) from wild-type mean generates a size-matched dataset for comparing wild-type and *Megf10* mutant soma enclosure (wild-type, n = 39 arbors; mutant; n = 33 arbors). (**J**) : Box and whisker plot (as in C) quantifying P0 Sox2^+^ cell enclosure in *Megf10* mutants and wild-type controls using the size-matched dataset (I). Mutant arbors enclosed more cells than wild-type arbors, consistent with impaired dendrite-soma exclusion in mutants. Mutant arbors also enclosed more somata than expected by chance (unmatched condition) suggesting a tendency towards soma aggregation in mutants. Sample sizes: real arbors as in I; unmatched images: wild-type, n = 38 per arbor, n = 1482 total; mutant, n = 32 per arbor, n = 1056 total. (**K**) Histogram of enclosure frequencies (as in D,E) using *Megf10* mutant arbors. Histogram for real mutant arbors (sample size as in I) was indistinguishable from chance rate measured using unmatched controls (n = 906; unmatched arbors enclosing zero cells were excluded). Statistics as in D,E. **(N**) Enclosed cell indices for individual P0 arbors were plotted (as in F) for the size-matched dataset (left) and the full dataset of mutant ON reference cells (right; see F for full wild-type dataset). Wild-type arbors enclosed fewer Sox2 cells than expected by chance (i.e., unmatched condition) but mutant arbors enclosed a similar number of neighboring cells in both real and unmatched conditions. Sample sizes: Left, as in J,K; right, n = 58 real arbors, 32 unmatched images per arbor (zero values excluded, n = 1,430 total controls). Error bars, mean ± SD (H,I); mean ± SEM (D,E,K) median ± 95% CI (F,L); min-max values (C,J). P-values shown on graphs; red text denotes value below alpha threshold. Scale bars, 10 µm. Also see Figure S2.

While the fundamental mechanism for exclusion zone formation appears quite similar for OFF and ON starburst cells, our analysis did uncover one notable difference between the two cell types: Whereas OFF starbursts become randomly positioned in *Megf10* mutants (Fig. 1), we observed that loss of *Megf10* unveils an attractive interaction among ON starburst cells. Two lines of evidence support this finding. First, ON *Megf10^−/−^* exclusion zone sizes were smaller, and mosaic regularity was lower, than would be expected for an array of randomly distributed cells (Fig 6C-E). This finding suggests that the mutant ON arrays are not in fact random, but instead display some mild aggregation behavior. We also observed a mild tendency toward aggregation in our analysis of soma enclosure by ON *Megf10^−/−^*Chat:mGFP^+^ arbors (Fig. 7J). Second, we noted anatomical examples of ON starburst clustering in *Megf10* mutants (Fig. 6A). Clumps were particularly notable at P15 in far peripheral retina (Fig. 8A), within regions that were typically outside the areas used for spatial statistical analysis shown in Fig. 6. By contrast, mutant OFF starburst arrays did not show evidence of attraction, either in anatomical images (Fig. 8A) or in the Chat:mGFP dendrite-soma analysis (Fig. 4L). Furthermore, mutant OFF starburst arrays are well modeled by a random point process (Fig. 1C-E). Together these results indicate that loss of MEGF10-mediated dendrite-soma exclusion has distinct effects upon the OFF and ON starburst arrays: Without MEGF10, OFF starbursts are randomly positioned whereas ON starbursts tend to attract each other.

**Figure 8:**
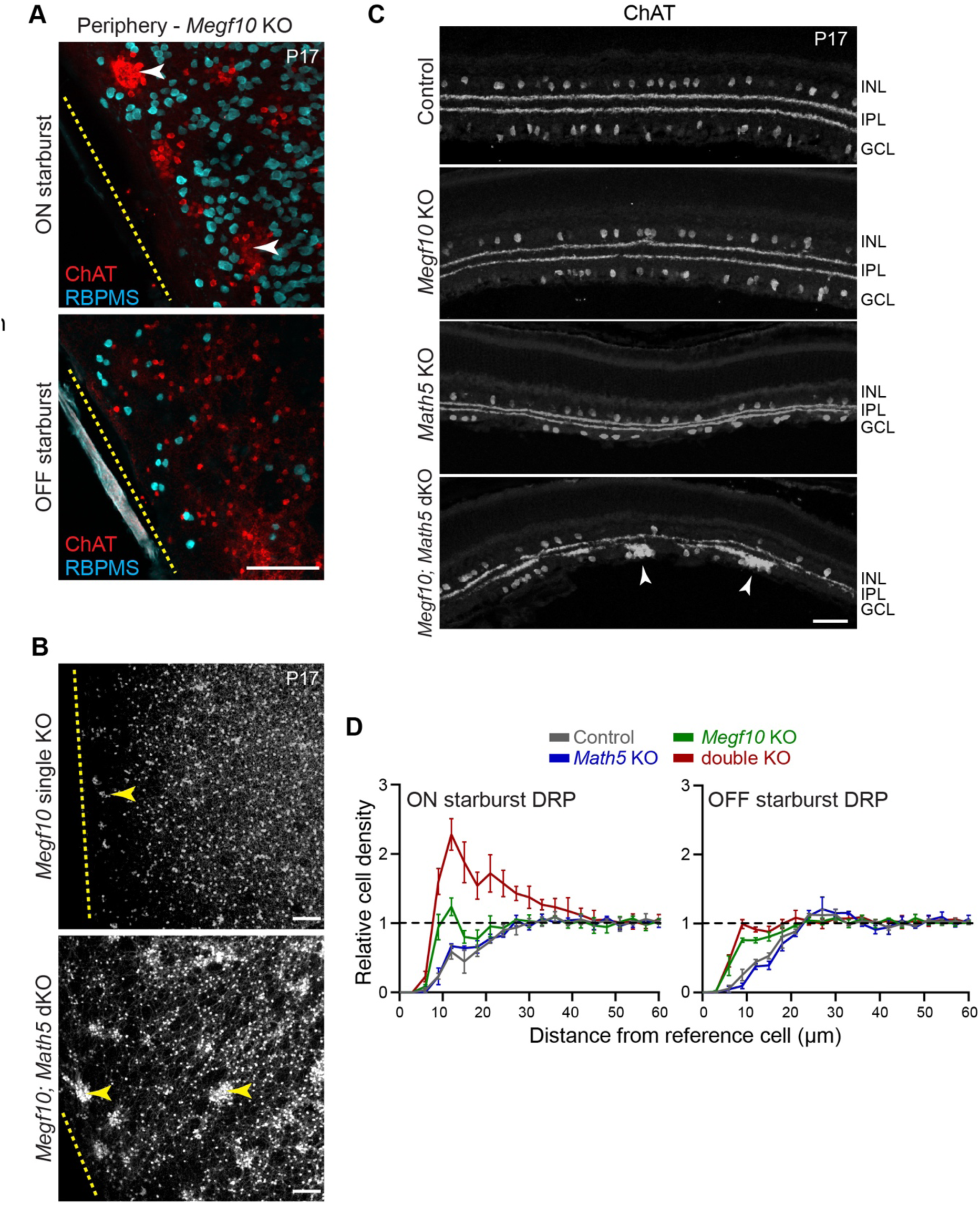
ON starburst aggregation in the absence of MEGF10 and retinal ganglion cells. (**A**) En face images of a representative *Megf10^−/−^* P17 retinal wholemount, showing ON starburst clumps at far retinal periphery. Images from single confocal stack at level of GCL (top) or INL (bottom). Dashed line, edge of retina. Anti-ChAT (red) labels starbursts; anti-RBPMS (blue) labels RGC somata. Mutant ON starbursts (top) form multicellular aggregates near retinal edge (arrows), while OFF starbursts do not (bottom). (**B**) Lower-magnification images encompassing larger region of peripheral retina than (A). Representative examples of ON starburst phenotype in ChAT-stained *Megf10* single knockout (KO) and *Megf10; Math5* double knockout (dKO). In single mutants (top), multicellular aggregates (arrow) are limited to the region directly adjacent to retinal edge (dashed line). In dKOs (bottom), clumping extends throughout imaged region, and clumps are larger than in single KOs. (**C**) P17 cross-sections through central retina, stained with anti-ChAT to reveal starburst cells and their IPL arbors. In controls (*Megf10^+/+^; Math5^+/+^*), as well as *Math5* and *Megf10* single KOs, few soma clumps are evident. Aggregates in *Megf10* single KOs are limited to ∼3 cell bodies. In *Megf10; Math5* dKOs (bottom), large clumps containing numerous ON starburst somata are seen within GCL (arrowhead). OFF starbursts in INL are unaffected. Images are representative of tissue from at least two different animals. (**D**) Density recovery profiles (DRPs) for P17 ON and OFF starburst arrays of each genotype. Annulus size, 3 µm. In wild-type and *Math5* single mutants, starburst density is lower at short intercellular distances than global cell density (dashed line), indicative of local cell-cell avoidance (i.e. exclusion zones). In dKOs, ON cell density (left) is higher at short spatial scales than global density, demonstrating cell aggregation. *Megf10* single KOs also show mild ON cell aggregation, but far less than dKOs. No such spike above global density is observed for OFF starbursts (right). Error bars, SD. Scale bars, 100 µm (A,B); 50 µm (C).

### Retinal ganglion cells influence ON starburst mosaic patterning

Finally, we investigated the origins of ON starburst clumping behavior in *Megf10* mutants. The incidence and size of abnormal starburst clumps was largest at the far retinal periphery (Fig. 8A), suggesting that the factors promoting homotypic attraction are most prevalent in peripheral retina. One crucial factor that varies along the center-peripheral axis is the density of retinal ganglion cells (RGCs), the largest cell population residing within the GCL alongside ON starburst cells. At the far periphery of mouse retina, RGC density is over two-fold lower than in central retina (Rodriguez et al., 2014; Wang et al., 2017). By contrast, the density of ON starburst cells declines only minimally (∼1.33-fold) between center and periphery (Keeley et al., 2007). Therefore, in the retinal region where clumping is most prominent, starburst cells comprise a larger fraction of GCL neurons than in regions where clumping is rare. This observation raises the possibility that heterotypic interactions with RGCs might shield ON starbursts from engaging in homotypic attractive interactions.

If this model is correct, experimental removal of RGCs should enhance the *Megf10^−/−^* starburst aggregation phenotype. To test this prediction we used *Math5* null mutant mice, in which RGC fate specification and survival defects lead to a near-complete absence of RGCs (Brodie-Kommit et al., 2021; Brown et al., 2001; Wang et al., 2001). The *Math5* mutant allele was crossed into the *Megf10* mutant background, thereby generating *Megf10^−/−^; Math5^−/−^* double knockout (dKO) mice. In dKO animals, the density of starburst neurons was similar to wild-type controls (OFF WT: 1495 ± 107.4 cells/mm^2^; OFF dKO: 1648 ± 102.8 cells/mm^2^; ON WT: 1290 ± 39.8 cells/mm^2^; ON dKO: 1310 ± 8.1 cells/mm^2^; mean ± SEM, n = 3 animals per genotype). However, spatial analysis using DRP revealed a major enhancement of ON starburst aggregation in dKOs relative to wild-type or to *Megf10* single mutants (Fig. 8D). This effect was specific to ON starbursts; OFF starbursts within the INL were not perturbed by removal of RGCs (Fig. 8D). In contrast to *Megf10* single mutants, in which large starburst clumps were restricted to the extreme periphery, we observed substantial clumping behavior throughout the dKO retina (Fig. 8B,C). Furthermore, the size of the starburst aggregates was typically much larger in dKOs than in *Megf10* single mutants (Fig. 8B). These results demonstrate that absence of RGCs increases the frequency of attractive interactions among ON starburst neurons. Importantly, starburst exclusion zones were normal in *Math5* single mutants (Fig. 8C,D). Thus, MEGF10-mediated repulsion is sufficient to overcome any enhanced attraction resulting from removal of RGCs. Together, these findings suggest that ON starbursts possess an intrinsic homotypic attractive activity, which is countered both by MEGF10 repulsion and by the presence of heterotypic neurons which serve to buffer starburst cells from interacting and adhering to each other.

## DISCUSSION

In this study, we identify a mechanism by which contact among homotypic retinal neurons leads to cell-cell repulsion and mosaic patterning. It has long been recognized that establishment of retinal mosaics involves dendritic contacts: As newly differentiating neurons begin growing dendrites and make first contact with their homotypic neighbors, a repulsive signal is initiated that leads to exclusion zone formation (Reese and Galli-Resta, 2002). Here we provide an explanation for how dendritic contacts create exclusion zones. Previously, the prevailing model was that homotypic contacts occur at dendritic tips, leading to dendrite tiling that carves out a unique territory for each cell (Cook and Chalupa, 2000; Huckfeldt et al., 2009; Reese and Keeley, 2015) (Fig. 3B). Because most retinal neurons do not tile their dendrites at maturity (Diamond, 2017; Reese and Keeley, 2015; Wässle and Riemann, 1978), a key prediction of this model is that mosaics are established during a transient period of dendritic tiling during early development (Huckfeldt et al., 2009). Critical tests of this model therefore require specialized anatomical tools: To assess transient tiling, one must generate strong and selective labeling of a single type of retinal neuron, including their nascent dendrites, from the earliest stages of their differentiation. Here we developed mouse genetic tools that enabled this type of anatomical investigation for starburst amacrine cells. We find that starburst dendrites never tile, but instead contact neighboring starburst somata immediately upon the initiation of dendrite outgrowth. These dendrite-soma contacts set up an anatomical arrangement in which neighboring starburst somata are excluded from each cell’s dendritic territory (Fig. 9A). Our analysis of *Megf10* mutants provides functional evidence that dendrite-soma exclusion is required for exclusion zone formation and mosaic patterning. Thus, we conclude that dendrite-soma contacts are well positioned to serve as the cellular mechanism for establishing starburst mosaics.

**Figure 9:**
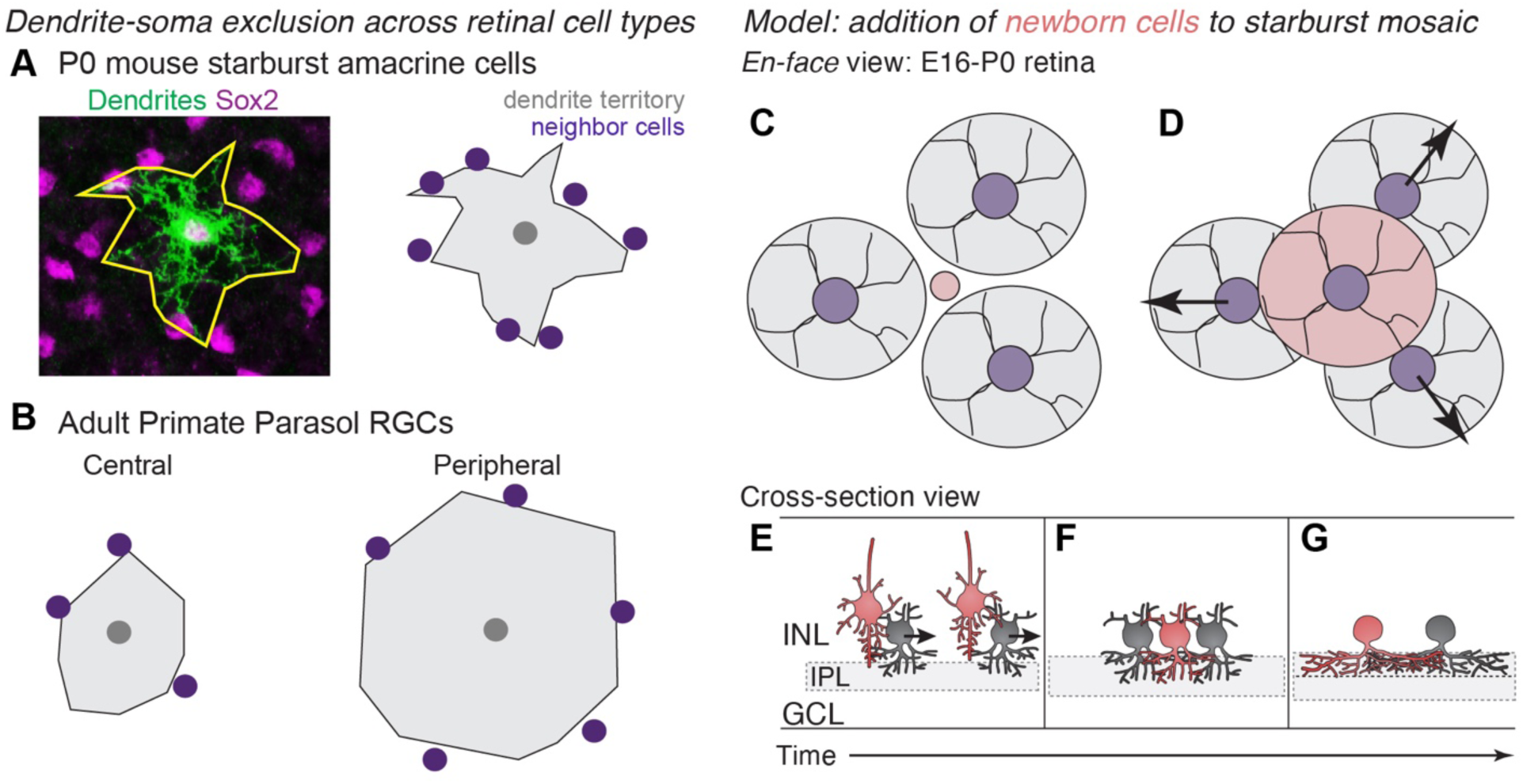
Summary and working model for mosaic patterning by homotypic dendrite-soma exclusion. (**A**) Summary of our findings: Newborn starburst neurons transiently establish dendritic territories that exclude homotypic neighbors, thereby generating local cell-cell repulsion that provides mosaic patterning. Left, photomicrograph showing P0 Chat:mGFP^+^ starburst cell dendrites (green; yellow outline) and locations of Sox2^+^ starburst neighbors (purple). Right, schematic drawing of dendritic territory and neighboring starburst cell body locations. (**B**) Parasol RGCs in adult macaque retina show precise alignment of dendrite boundaries with homotypic somata, resembling our starburst findings (A); this suggests other cells may also use dendrite-soma exclusion to achieve mosaic patterning. Drawing summarizes results of Dacey and Brace (1993). Colors as in A. Other retinal cell types also show this pattern; see Discussion. (**C-G**) Working model for how newborn starburst cells use dendrite-soma exclusion to integrate into the developing mosaic. Illustrations show this process from en face (C,D) or cross-sectional (E-G) viewpoint. C, A radially migrating newborn starburst cell (red) extends its leading process into the nascent INL. Earlier-born starbursts (gray) are already regularly spaced (Galli-Resta et al, 1997). The red cell then initiates dendrite morphogenesis, generating new arbors that contact homotypic neighbors on their somata (D,E), leading them to move tangentially to make room for the newly-arrived cell (arrows). Because these tangential movements are limited by the presence of neighboring dendrites, which prevent entry of the tangentially migrating cell, a regularly spaced network is established whereby dendrite-soma contact determines exclusion zone size (D,F,G). Initial dendrite-soma contacts occur within INL or GCL, using transient soma layer arbors (E,F), but subsequent contacts use IPL arbors which contact the base of the neighboring soma to establish an exclusion zone (G). Drawing in C,D inspired by Cook and Chalupa (2000).

Based on our present results, as well as prior studies showing that regular spacing is maintained throughout the period when starbursts are generated (Galli-Resta et al., 1997), we propose the following working model for starburst mosaic patterning (Fig. 9C-G). As a newborn starburst neuron arrives at the inner retina from the outer neuroblast layer (Fig. 9C,E), it first engages in arbor growth that is directed towards the somata of other starbursts that arrived earlier (Fig. 3F). This behavior generates a transient network of starburst dendro-somatic homotypic contacts within the INL and GCL (Ray et al., 2018; Fig. 3D,E; see model, Fig. 9E,F). At this stage, starbursts remain motile in the tangential plane (Reese et al., 1999), but their movements are limited by the location of their homotypic neighbors’ dendritic territories. Thus, regular spacing begins to emerge as each cell settles at a location where it does not overlap a neighboring dendritic arbor (Fig. 9C-F). Subsequently, the newborn cell begins to ramify dendrites within the IPL, which terminate at the base of neighboring starburst cell bodies and occasionally branch out into the soma layer for more extensive cell body contact (Fig. 3J,K; see model, Fig. 9F,G). Like the transient soma-layer arbors, these IPL arbors also serve to exclude homotypic cell bodies. Thus, growth of the newborn cell’s arbors leads the pre-existing cells to adjust their positioning to incorporate the new cell (arrows, Fig 9D,E). In this way, the newborn starburst cell carves out an exclusion zone for itself that is defined by the extent of its dendritic arborization (Fig. 9D,G).

There is precedent for the idea that regular spacing of retinal neurons could involve exclusion of homotypic somata from dendrite territories. In the adult primate retina, parasol cells exhibit dendrite-soma arrangements that are strikingly similar to the arrangements of developing starburst neurons (Fig. 9A,B): Parasol somata in the GCL are precisely aligned over the edges of their neighbors’ IPL dendritic territories (Dacey and Brace, 1992). This alignment is maintained despite large variations in cell density and dendritic size across different retinal eccentricities (Dacey and Brace, 1992), which suggests that alignment results from dendrite-soma coordination during development. Similarly, mouse and primate horizontal cells also grow dendrites that extend as far as the neighboring cell body; and they also show coordination between arbor size and soma positioning when cell numbers are varied (Reese and Keeley, 2015; Wässle et al., 2000). In addition to these two well-documented examples, dendrite-soma alignment can be appreciated for numerous mouse RGC types within the online “RGC museum” (Bae et al., 2018); and for mouse Vglut3^+^ amacrine cells (Keeley et al., 2021). Further anatomical analysis will be needed to show that these latter cell types indeed exhibit selective dendrite-soma alignment; however, these observations raise the possibility that such alignments may be a widespread phenomenon. In this case, dendrite-soma exclusion could be involved in the mosaic patterning of many retinal cell types beyond starburst neurons. Dendrite-soma exclusion is only a transient developmental phenomenon for starburst cells (Fig. 3A), but they may be unusual in this respect, as mature starbursts exhibit an exceptionally high dendritic overlap factor (Keeley et al., 2007). Other cell types, which do not require so much overlap, may simply cease growing their dendrites once dendrite-soma exclusion has been established, thereby allowing the exclusionary relationship to be appreciated even in mature retina (Fig. 9B).

Key support for the dendrite-soma exclusion model comes from our analysis of *Megf10* mutants, which lack starburst exclusion zones from the earliest age at which we could measure them (P0; Fig. 1F, Fig. 6D). Therefore, whatever cellular mechanism establishes exclusion zones must be severely compromised in these mutants. Here we show that *Megf10* mutant starburst cells are unable to exclude their homotypic neighbors from their dendritic territories, strongly implicating this repulsive behavior in exclusion zone formation. Why are mutant neurons unable to accomplish this exclusion? If starburst somata were immotile, this would render any repulsive signal from dendrites ineffective; however, we found that mutant starbursts move just as frequently and just as far as wild-type cells, ruling out this possibility (Fig. 5). The fact that *Megf10* mutant starbursts have smaller dendrite arbors at P0 (Fig. 4J; Fig. 7H) may be one factor contributing to their failure to exclude neighbors: With smaller arbors, mutant cells are less likely to contact neighboring somata to initiate repulsion. However, this phenomenon cannot fully explain the *Megf10* phenotype, because when we controlled for arbor size in our anatomical studies we found that *Megf10* mutant arbors are still defective at excluding neighboring somata. Moreover, loss of *Megf10* only diminishes IPL arbors, whereas transient soma-directed arbors in the INL and GCL are apparently normal or even increased (Ray et al., 2018), thus providing ample opportunities for neighboring cells to interact in mutants (Fig. 3E). Therefore, a lack of contact between neighboring cells does not suffice to explain the mutant phenotype. Instead, we favor the idea that cytoskeletal properties of mutant arbors may be affected in a manner that diminishes their ability to resist entry by neighboring cells. This is plausible because MEGF10 and its invertebrate homologs, Draper and CED-1, are known regulators of the actin cytoskeleton (Kinchen et al., 2005; Williamson and Vale, 2018); however, further experiments will be needed to test this model.

Our studies of ON starburst cells highlight a second type of cellular mechanism that influences mosaic patterning. In addition to homotypic dendrite-soma exclusion, ON starbursts also engage in heterotypic interactions with RGCs that insulate them from excessive homotypic clumping. The involvement of heterotypic interactions is surprising, because mosaic patterning was previously considered to arise mainly through homotypic interactions. Besides ON starbursts, several other retinal cell types are reported to exhibit homotypic adhesion/attraction which is countered through the action of the repulsive cell-surface molecules (Fuerst et al., 2009; Fuerst et al., 2008; Keeley et al., 2011). In these other cell types, mosaic disruptions due to cell clumping can be induced by removing the relevant repulsive molecules (DSCAM or DSCAML1), or by increasing the number of neurons via genetic manipulations that prevent developmental apoptosis (Reese and Keeley, 2015). Here we find that removing neighboring GCL cells can also lead to ON starburst clumping. While *Megf10*-mediated homotypic repulsion needs to be compromised in order to reveal the full extent of this effect, even in wild-type retina ON starbursts are more prone to clumping than OFF starbursts (Galli-Resta, 2000). This finding suggests that inadvertent homotypic adhesion is more likely in the GCL, which is a monolayer and has a much higher overall fraction of starburst neurons than the INL. Taken together, these findings support the idea that increasing the likelihood of close homotypic contacts – whether through increases in cell numbers or removal of insulating neighbors – can counter molecular repulsion mechanisms and in some cases overwhelm them, leading to homotypic neuron aggregation.

While we did not find evidence for dendritic tiling among developing starburst neurons, tiling may still have a role in exclusion zone formation for other cell types. For example, midget RGCs in primate retina have tiled dendritic arbors, with territory sizes that scale inversely with cell density across eccentricities (Dacey, 1993). This observation raises the possibility that the two major types of primate RGCs establish mosaic patterning in different ways – midget cells via tiling, and parasol cells via dendrite-soma exclusion (Fig. 9B). Another example is mouse horizontal cells, which generate transient vertically-oriented arbors that tile during the perinatal period (Huckfeldt et al., 2009). However, immature horizontal cells also generate dynamic lateral arbors that overlap instead of tiling – and may in some cases contact neighboring somata (Barrasso et al., 2018). Furthermore, as noted above, their mature dendrite arrangements are consistent with the idea that dendrite-soma exclusion could also influence horizontal cell mosaic patterning (Reese and Keeley, 2015). Future studies taking advantage of *Megf10; Megf11* double mutants, which lack horizontal cell exclusion zones (Kay et al., 2012), may be able to establish the relative importance of each mechanism.

Altogether, our study highlights new cellular mechanisms that contribute to mosaic patterning for ON and OFF starburst cells, and which have the potential to pattern many other cell types as well. It will be important to conduct future studies understand the extent to which this mechanism, as opposed to tiling or other modes of homotypic interaction, may explain exclusion zone formation by other retinal cell types.

## ACKNOWLEDGEMENTS

This work was supported by grants from the National Eye Institute (EY024694 and EY031445 to JNK; EY005722 to Duke University) and from Research to Prevent Blindness (Unrestricted grant to Duke University). We thank Gary Kucera and Cheryl Bock (Duke Transgenic Mouse Shared Resource) for generating the *Megf10^CreNeo^* mice; Stephanie Fogerson for assistance with microscopy; Vanessa Puñal for providing MATLAB and Fiji scripts; Ari Pereira for technical assistance and animal colony management; Joshua Winer, Brigid Hogan, and Tom Glaser for providing mice; Marla Feller and John Pearson for helpful discussions; and Caitlin Paisley and the rest of the Kay lab their feedback on the manuscript.

## MATERIALS AND METHODS

### Animals

Experiments conducted on animals were reviewed and approved by the Institutional Animal Care and Use Committee of Duke University. The animals were maintained under a 12 hour light-dark cycle with *ad lib* access to food and water. Animals of both sexes were used in this study. Wildtype CD1 mice were purchased from Charles River Laboratories.

For these studies, we used several existing transgenic and mutant mouse lines: (1) *Megf10^tm1b(KOMP)Jrs^*, referred to as *Megf10^−^* or *Megf10* KO, in which the *Megf10* locus drives expression of a *lacZ* reporter instead of the endogenous MEGF10 protein (Kay et al., 2012); (2) *Chat^tm2(cre)Lowl^* (RRID:IMSR_JAX:006410), referred to as *Chat^Cre^* (Rossi et al., 2011); (3) A germline Flp deleter strain, Tg(ACTFLPe)9205Dym/J (RRID:IMSR_JAX:005703); (4) Tg(Pax2-cre)1Akg/Mmnc (RRID:MMRRC_010569-UNC), known as Pax2-Cre (Ohyama and Groves, 2004); (5) *Atoh7^tm1Gla^*(RRID:MMRRC_042298-UCD), also known as *Math5^−^* or *Math5* KO (Brown et al., 2001). We additionally used three different Cre reporter strains. Two of these strains express membrane-targeted green fluorescent protein (mGFP) upon Cre recombination: (1) *Gt(ROSA)26^Sortm4(ACTB-tdTomato,-EGFP)Luo^* (RRID:IMSR_JAX:007576), also known as mT/mG (Muzumdar et al., 2007); (2) Gt(ROSA)26Sor^tm1(CAG-EGFP)Blh^ (MGI:3850169), also known as *Rosa26^fGFP^* (Rawlins et al., 2009). The third Cre reporter strain expresses tdTomato fluorescent protein upon Cre recombination: *Gt(ROSA)26Sor^tm14(CAG-tdTomato)Hze^* (RRID:IMSR_JAX:007914) (Madisen et al., 2010).

Mouse lines were obtained from Jackson Laboratories except the *Megf10* null mutant strain, which we previously generated; the *Math5* null mutant strain (kind gift of Tom Glaser, UC Davis); the *Rosa26^fGFP^* strain (kind gift of Brigid Hogan, Duke University); and the Pax2-Cre strain (kind gift of Joshua Weiner, University of Iowa). The *Tm1a* allele of *Megf10*, which can be converted to the null allele using a germline Cre deleter strain, is available from the Mutant Mouse Regional Resource Center (cat# MMRRC:068040-UNC).

### Generation of *Megf10* driven Cre mouse lines

Production of *Megf1^Cre^* knock-in mouse lines were generated in collaboration with Duke Transgenic Core. This genomic location was targeted using homologous recombination and a modified pL253 vector (Liu et al., 2003) with mc1-driven thymidine kinase for ES cell selection. Targeted ES cell clones were validated by PCR and southern blot analysis before embryo injection. Founder mouse lines were validated by PCR. These mouse lines were engineered to modify exon 25 of Megf10, replacing the endogenous stop codon with a FLAG-epitope tag, followed by a T2A self-cleaving peptide (Liu et al., 2017), to release a Myc-epitope tagged Cre recombinase. The founder mouse line, *Megf10^CreNeo^*, contained an FRT flanked neomycin resistance sequence downstream of the Cre. To generate the *Megf10^Cre^*allele, the neomycin cassette was removed by crossing *Megf10^CreNeo^*with a mouse line expressing germline FLP recombinase (see Animals section above for strain details). Removal of the neomycin resistance cassette was confirmed by PCR. During the coronavirus pandemic of 2020, due to severe constraints on mouse husbandry and staffing, we lost several mouse strains from our colony including the parental *Megf10^CreNeo^* strain. However, the *Megf10^Cre^* strain was not lost, and the targeting vector is still available in case the parental strain needs to be recreated.

### Mosaic analysis

Using confocal microscopy, 40x images (353.55 x 353.55 μm) were acquired from whole-mount retinal preparations in mid-peripheral retina. Z-stacks encompassed most of the inner retina, from the vitreal retinal surface to the middle of the INL. OFF and ON starburst populations were identified at different optical planes of these Z-stacks based on cell type-specific marker expression and their location in INL or GCL. Hoeschst counterstaining allowed identification of retinal layers within the Z-stacks. For mosaic analysis of developing OFF and ON starbursts (P0 and P5) we used anti-Sox2 as a cell type-specific nuclear marker (Whitney et al., 2014). Sox2 is also expressed by nerve fiber layer astrocytes and Müller glia, but these could easily be distinguished from starburst neurons based on laminar location and/or nuclear morphology. For later timepoints (i.e. P19), anti-ChAT was used to label ON and OFF starburst populations.

For measurements of cell size at P0 and P5, starbursts were labeled using antibodies to β-galactosidase in *Megf10^lacZ^* reporter mice (Kay et al., 2012; Ray et al., 2018). This marker was preferable to Sox2 for size measurements because it filled the cytoplasm. At P19, ChAT was used to measure cell size as described previously (Kay et al., 2012). Sizes were measured in ImageJ by encircling cells with an ROI; the Feret’s diameter tool was then used to measure the maximum diameter of the ROI. Sample sizes for diameter measurements were at least n = 100 cells from at least two animals. At all ages, mean soma size was 10.0 µm so we used this value for all downstream analysis steps.

For analysis of spatial statistics, images were loaded into FIJI/ImageJ software (Schindelin et al., 2012) and the point selector tool was used to manually mark the center of each starburst cell, generating X-Y coordinates. These coordinates were then used to produce Voronoi Domain Regularity Indices using Fiji software, and Density Recovery Profiles using WinDRP software as previously described (Kay et al., 2012; Rockhill et al., 2000) (see Supplemental Fig. S1 for an illustration of these methods). Exclusion zone sizes were calculated from the DRP in WinDRP software, which implements Rodieck’s definition of the effective radius (i.e. exclusion zone) as the midpoint of the rising part of the DRP curve (Rodieck, 1991). Annulus size increment for DRP analysis was 5 µm.

Random simulations were generated using a custom MATLAB script as previously described (O’Sullivan et al., 2017). Briefly, the algorithm placed cells into a square field of view matching the size of the ones used for real images (i.e. 353.55 µm on all sides). The cells were placed one-by-one according to a Poisson point process, until the density of the array was equal to the mean starburst density measured from real data (Fig. 1C). The only constraint on cell location was that two cells could not occupy the same physical location. To determine if two cells overlapped, each cell was assigned a diameter of 10 µm (the average cell size measured for real starburst neurons at all ages analyzed – see measurement details above). If a new cell was added to the array at a location where its diameter overlapped with a pre-existing cell, placement at that location was cancelled and a new location was assigned based on the Poisson point process.

For experiments examining *Math5; Megf10* double knockout retinas, mosaic analysis was performed using 20x images (636.5 x 636.5µm) and the annulus step size used for DRP analysis was 3 µm.

### Arbor territory analysis

The relationship between starburst dendritic arbor territories and neighboring homotypic somata was assessed using two channel images that contained 1) individually labeled starburst arbors from *Chat^Cre^*; mGFP reporter mice; and 2) co-staining with anti-Sox2 to label the nuclei of the entire ON and OFF starburst populations. Z-stacks encompassing the entire dendritic arbor and relevant nuclear layer were collected using confocal microscopy (Olympus FV300 or FV3000). Dendrite territories were drawn in ImageJ by lines connecting arbor tips to generate a series of polygonal regions of interest (ROIs) for each mGFP^+^ starburst in the dataset. We counted the number of Sox2^+^ homotypic somata completely contained within those territories, including the soma belonging to the reference cell.

To assess if starburst arbor territories contained fewer homotypic somata than expected by chance, we performed these dendritic Sox2 counts on two groups of images. First, as the experimental group, we used real images, in which the dendritic ROI was overlaid over the actual accompanying Sox2 channel. Second, as a control in which the relationship between dendrite area and neighboring soma position was severed, we used “unmatched” shuffled images in which the dendritic ROI was placed over non-matching Sox2 images. These unmatched control images were generated by transposing the dendritic ROI onto the Sox2 channel from other fields of view from the same dataset – i.e., dendrite polygons from wild-type ON starbursts were transposed onto wild-type ON Sox2 images, while *Megf10* mutant OFF polygons were transposed onto mutant OFF Sox2 images. Typically, each arbor was placed onto the Sox2 array of all other images from the same dataset, although there were rare instances where a certain Sox2 array was not used. This procedure generated a family of unmatched control images for each dendritic arbor (n = 17-38 simulations per arbor, varying based on the number of real images in each dataset; see figure legends for precise numbers). For some analyses (Supplementary Fig. S2) we also generated “mutant on wild-type” unmatched controls in which *Megf10* mutant polygons were transposed onto wild-type Sox2 arrays. After generating the unmatched control images, the number of Sox2 cells fully enclosed by the arbor polygon was manually counted in each simulation, using the methodology noted above for the real images. We then calculated unmatched enclosure rates for each individual arbor by averaging across the family of simulations generated from that arbor.

The unmatched dataset was used to test whether the frequency of Sox2 soma enclosure observed in the real data is lower than the frequency expected by chance. Box and whisker plots comparing enclosure numbers for real and unmatched arbors (e.g. Figs. 4C, 7C) were generated using the full simulation dataset. The individual arbor unmatched averages were used for two analyses: 1) frequency distribution histograms, which plotted the frequency with which *n* somata were enclosed by an individual arbor (e.g. Figs. 4D,E, 7D,E); and 2) calculation of the enclosed cell index for each arbor (e.g. Figs. 4F, 7F). These two analyses are described below in further detail. For these two analyses we excluded unmatched images that did not enclose any Sox2 cells; this was necessary because real images always contained at least one Sox2 cell – i.e. the soma of the reference mGFP^+^ cell. Therefore, without excluding zero values, the unmatched datasets were systematically skewed towards smaller values. Moreover, without this exclusion, chi-squared tests comparing enclosed cell proportions from real and unmatched datasets (see below) would not be valid because the “0” bin would differ between these groups for artificial reasons. Notably, even with the skew towards smaller values introduced by inclusion of zeroes in the unmatched datasets, we still found more enclosed cells in the unmatched groups (Figs. 4C, 7C).

#### Frequency distribution histograms

The “real” curves in these histograms represent the fraction of arbors that enclosed exactly *n* Sox2^+^ somata within the real images. To obtain the “unmatched” curves, we began by calculating the frequency with which each individual dendritic arbor ROI enclosed *n* Sox2^+^ somata across all of the unmatched control images using that arbor. This table of per-arbor enclosure frequencies was then used to calculate the mean (± SEM) frequency of enclosing *n* somata across all arbors in a given dataset, which is the value plotted in the figures (e.g. Fig. 4D,E). Chi-squared tests were used to test for differences between real and unmatched curves.

#### Enclosed cell index

To compare real Sox2 cell enclosure rates to chance rates at the individual arbor level, we calculated the enclosed cell index for each mGFP^+^ arbor. This was defined as:

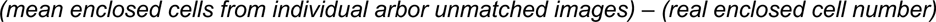

This index will be equal to zero if the number of cells enclosed by a given arbor in its real and unmatched images is the same. Deviations from zero indicate more cells enclosed by either real or unmatched arbors (positive = more in unmatched; negative = more in real). To evaluate deviations from zero, the index was plotted for each cell, and the median value was compared to zero using a Wilcoxon one-sample test. This procedure is tantamount to performing a Wilcoxon matched-pairs test comparing the number of Sox2 cells enclosed by real arbors vs. their unmatched controls.

### Analysis of dendro-somatic starburst contacts in the INL

Confocal en-face images of Chat:mGFP^+^ OFF starburst cells (P0-P1) were acquired from mice that also carried a single copy of the *Megf10^lacZ^*allele, such that the full starburst array could be revealed using antibodies to β-galactosidase (βgal). Using Z-stacks encompassing the full INL arbor of the GFP^+^ reference cell, the trajectory and termination site of each GFP^+^ arbor was examined in three dimensions, using 3D reconstructions and orthogonal views, as necessary, to score it as to whether it terminated upon a neighboring βgal^+^ cell. As a negative control measuring the frequency of soma contact that may be expected by chance, the same analysis was performed on Z-stacks wherein the GFP channel was flipped about the horizontal and vertical axes.

### Histology and immunohistochemistry

#### Retinal cross sections

Mice were anesthetized by hypothermia (P0 and P5 mice) or isoflurane (all other ages) and euthanized by decapitation. Eyes were enucleated, washed in PBS, and fixed in PBS containing 4% formaldehyde (pH 7.5) for 1.5 hours at 4°C. After fixation, eyes were washed with PBS (2x) and stored in PBS containing 0.02% sodium azide at 4°C until further processing. The eyecup was isolated and sunk in PBS containing 30% sucrose, then embedded in Tissue Freezing Medium (VWR) before being frozen in 2-methylbutane chilled by dry ice. Tissue sections were cut on a cryostat to 20 µm and mounted on Superfrost Plus slides and dried on a slide warmer. For antibody labeling, slides were washed for 5 min with gentle agitation in PBS to remove embedding medium and blocked for 1 hr in PBS + 0.3% Triton X-100 (PBS-Tx) containing 3% normal donkey serum. Primary antibodies were diluted in blocking buffer, added to slides, then incubated overnight at 4° C. Slides were washed with PBS 3X for 10 min followed by incubation with secondary antibodies diluted in PBS-Tx for 1–2 hr at RT. Slides were washed again with PBS 3X for 10 min then coverslipped using Fluoromount G.

#### Retinal whole-mounts

After obtaining eyecups as described above, retinas were dissected free of the eyecup, washed in PBS, then blocked for 3 hours with agitation at 4° C in blocking buffer (constituted as described above). Primary antibodies were diluted in blocking buffer, added to retinas, and incubated for between 5 and 7 days on a rocker at 4°C. After primary staining, retinas were washed 3 times with PBS over 2 hours and then incubated in secondary antibodies (diluted in PBS-Tx). Secondary staining was performed overnight at 4° C with gentle agitation. Retinas were again washed in PBS (3x) over 2 hours on a room temperature rocker. For mounting on slides, four 90° radial incisions were made, approximately 1/3 the radius of the retina. Retinas were then flattened on nitrocellulose paper (Millipore) photoreceptor side down and coverslipped with Fluoromount G (Southern Biotech).

### Statistical analysis

All analyses were performed in JMP 12 (SAS Institute) or GraphPad Prism 10. Parameters of statistics (i.e. sample size, tests conducted and P-values) can be found within figures or figure legends. For analysis of Sox2 cell enclosure by starburst dendritic arbors, nonparametric statistical tests (Mann-Whitney U test, Wilcoxon test) were used because of the non-Gaussian distribution of values (see for example Fig. 4C, Fig. 7C).

---

**Supplemental Fig. S1:**
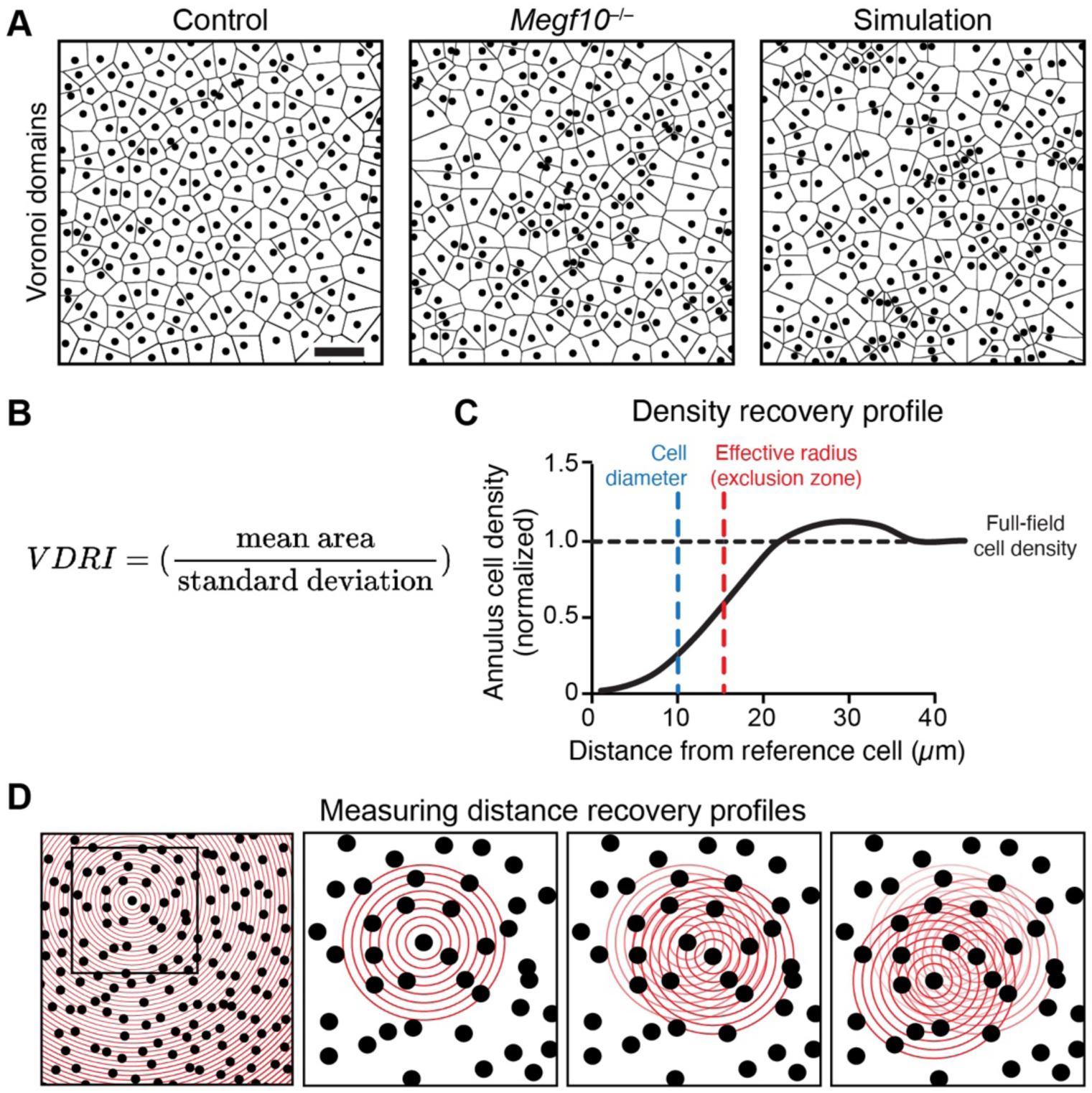
Methods used for analysis of starburst spatial patterning. (**A,B**) Quantification of mosaic regularity using Voronoi domain regularity analysis. (**A**) Representative examples of OFF starburst soma locations (circles) and their Voronoi domains from P5 wild-type retina (left); P5 *Megf10* mutant retina (center); or a computer-generated random simulation (right). Circles represent cell locations and are sized to approximate the average diameter of P5 OFF starburst somata. Voronoi domains, defined as the set of points closest to any given cell, are indicated by lines/polygons. For left and center panels, cell locations were derived from real images like those in Fig. 1B. Random simulations (right) were matched to P5 cell size and density measured from real images (see Methods). In wild-type arrays, the areas of the Voronoi domains are fairly uniform, reflecting the uniform distribution of starburst neurons in wild-type retina. By contrast, in *Megf10* mutants and in random simulations, Voronoi areas are more variable in size. (**B**) Formula for computing Voronoi domain regularity index (VDRI): The mean Voronoi area for a given image is computed and divided by the standard deviation of the area sizes. Note that Voronoi domains touching the edge of the field of view are excluded from the analysis. **(C,D**) Quantification of exclusion zone sizes using the density recovery profile (DRP). (**C**) Schematic showing key features of the DRP plot. The plot shows how the density of cells within annuli of *d* distance from any given cell compares to the overall cell density within an image (horizontal dashed line). (**D**) Illustration of the procedure used to compute the DRP. For each cell in the array, a series of rings of increasing size (step size used here, 3-5 µm) are drawn with the index cell at the center. The density of cells lying within each annulus is computed. After the procedure is repeated for each cell in the array, the average cell density for each annulus is calculated, normalized to the overall cell density in the entire image, and plotted as in C. In the example graph (C), which is a schematic illustrating typical starburst measurements, rings close to the reference cell have lower cell density than the overall image, indicating short-range cell-cell repulsion that prevents homotypic neighbors from settling near each other. At longer distances, annulus cell density approximates the overall cell density. The effective radius (i.e. exclusion zone) is measured as the midway point of the rising portion of the curve (red vertical dashed line). If cells are randomly positioned – i.e. they do not show local cell-cell repulsion or attraction – then the only constraint on their position is that two cells cannot occupy the same physical location in the 2D plane. As such, the exclusion zone size measured by DRP will be approximately equal to the average cell diameter (blue line). In our analysis of OFF starbursts, both real *Megf10* mutant arrays and random simulated starburst arrays (e.g. A, center & right panels) had measured exclusion zones similar to the average starburst cell diameter (Fig. 1). For cell types that form mosaics, such as wild-type starbursts, exclusion zones are larger than the cell diameter (compare vertical dashed lines), which indicates presence of *bona fide* cell-cell avoidance. The DRP can also be used to measure cell-cell attraction, which is indicated by the presence of annuli with a higher cell density than the overall image (i.e. above the horizontal dashed line, not shown here but see Fig. 8). In these cases, cells may be pulled out of their 2D plane into a clump, which can lead to exclusion zone size measurements that are lower than the average cell diameter (as cells will appear to overlap in the photomicrographs used for DRP analysis). This phenomenon was observed for ON starbursts in *Megf10* mutants (Fig. 6).

**Supplemental Figure S2:**
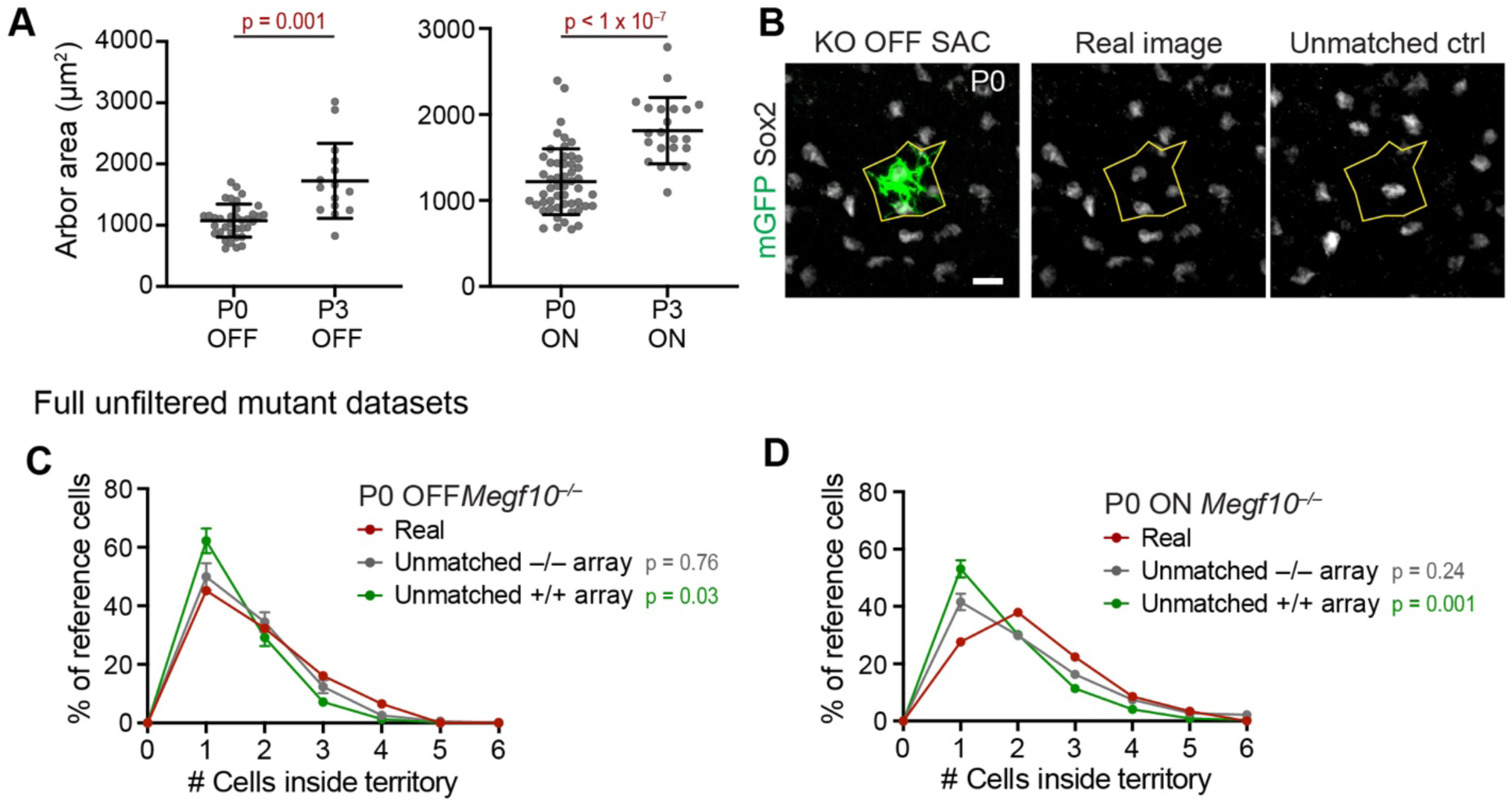
Additional analysis of starburst dendritic arbor size and soma exclusion. (**A**) Wild-type starburst arbors are larger on average at P3 than at P0. Left, OFF starbursts; right, ON starbursts. Statistics: two tailed t-test, p-values shown on graph. Sample sizes (OFF): P0, n = 36; P3, n = 15. Sample sizes (ON): P0, n = 52; P3, n = 22. Error bars, mean ± SD. (**B**) Example image showing a *Megf10^−/−^* OFF starburst with a small dendritic territory; this cell was outside the 1-standard-deviation cutoff used for the analysis in Figure 4. Left, center panels show real image; right panel shows unmatched control image in which the dendritic polygon was placed onto a mutant OFF starburst array from a different retinal location. Sox2 marks neighboring starburst somata. Scale bar, 10 µm. (**C,D**) Frequency distribution histograms showing Sox2^+^ cell enclosure for the full unfiltered *Megf10^−/−^* OFF (C) or ON (D) datasets. To measure enclosure frequencies expected by chance, we generated two types of unmatched controls: one in which real mutant arbor polygons were placed onto mutant (–/–) Sox2 arrays (gray; this was the methodology used for Fig. 4M and Fig. 7K), and one in which these polygons were placed onto wild-type (+/+) Sox2 arrays (green). Real and mutant-unmatched curves were not statistically distinguishable, consistent with a lack of dendrite-soma exclusion in mutants. This finding is in accord with our results using size-matched filtered datasets (Figs. 4M and 7K). If the +/+ unmatched curve is taken as the chance enclosure rate, it would appear that more cells are enclosed by real mutant arbors than expected by chance. This result may suggest possible starburst aggregation in mutants, or it may mean that using the mutant Sox2 array provides a more accurate measure of the chance enclosure rate. However, it is not consistent with presence of dendrite-soma exclusion, which would have caused the real mutant curve to be shifted left relative to the control distribution. Thus, regardless of whether the filtered or full datasets are used, and regardless of whether mutant unmatched simulations are generated using mutant or wild-type Sox2 arrays, our conclusion that dendrite-soma exclusion is absent in mutants remains unchanged. Sample sizes: mutant OFF n = 29 real arbors, 20 unmatched images per arbor; mutant ON n = 58, 32 unmatched images per arbor. Since “zero enclosed cells” is an impossible value for the real data (due to the presence of the reference cell body), we excluded zeroes counted from unmatched simulations (final sample sizes: OFF –/– unmatched, n = 479; OFF +/+ unmatched, n = 777; ON –/– unmatched, n = 1,449; ON +/+ unmatched, n = 1,786). Statistics, chi-squared tests comparing the real distribution to each of the unmatched control distributions. P-values shown on graph. Error bars, mean ± SEM.

